# SyNPL: Synthetic Notch pluripotent cell lines to monitor and manipulate cell interactions *in vitro* and *in vivo*

**DOI:** 10.1101/2021.09.24.461672

**Authors:** Mattias Malaguti, Rosa Portero Migueles, Jennifer Annoh, Daina Sadurska, Guillaume Blin, Sally Lowell

## Abstract

Cell-cell interactions govern differentiation and cell competition in pluripotent cells during early development, but the investigation of such processes is hindered by a lack of efficient analysis tools. Here we introduce SyNPL: clonal pluripotent stem cell lines which employ optimised Synthetic Notch (SynNotch) technology to report cell-cell interactions between engineered “sender” and “receiver” cells in cultured pluripotent cells and chimaeric mouse embryos. A modular design makes it straightforward to adapt the system for programming differentiation decisions non-cell-autonomously in receiver cells in response to direct contact with sender cells. We demonstrate the utility of this system by enforcing neuronal differentiation at the boundary between two cell populations. In summary, we provide a new tool which could be used to identify cell interactions and to profile changes in gene or protein expression that result from direct cell-cell contact with defined cell populations in culture and in early embryos, and which can be adapted to generate synthetic patterning of cell fate decisions.

**SUMMARY STATEMENT:** Optimised Synthetic Notch circuitry in mouse pluripotent stem cells provides a modular tool to monitor cell-cell interactions and program synthetic patterning of cell fates in culture and in embryos.

## INTRODUCTION

During embryogenesis, pluripotent cells undergo a series of cell fate decisions that are controlled by interactions between epiblast cells, their early differentiated derivatives, and the surrounding extraembryonic tissues (Arnold and Robertson, 2009; Nowotschin and Hadjantonakis, 2010; Rossant and Tam, 2009). The transcriptional changes that accompany exit from pluripotency and differentiation into specific cell types have been extensively characterised, and the long-range signals that control these changes are now well understood (De Los Angeles et al., 2015; Kinoshita and Smith, 2018; Pera and Tam, 2010; Posfai et al., 2021; Tam and Loebel, 2007). Less is known about how early developmental decisions are influenced by direct interactions of cells with their neighbours. Cell-cell interactions play a key role in development (Dias et al., 2014; Gurdon, 1987; Johnson and Ziomek, 1983; Schultz, 1985), but until recently there has been a paucity of molecular and technological tools available to study these processes in detail in relevant settings (Nishida-Aoki and Gujral, 2019; Yang et al., 2021).

Quantitative image analysis can be used to identify and infer the effect of neighbours on the properties of cells of interest in fixed samples (Blin et al., 2018; Fischer et al., 2020; Forsyth et al., 2021; Toth et al., 2018). We have recently developed a software suite for automated neighbour identification during live imaging (Blin et al., 2019), which provides researchers with a further dimension to study the effects on cell-cell interactions on cell fate decisions. Whilst live image analysis provides high-resolution visual information, this approach is labour-intensive and only leads to neighbour identification *a posteriori*.

The field of Synthetic Developmental Biology (Davies, 2017; Ebrahimkhani and Ebisuya, 2019; Ho and Morsut, 2021; Santorelli et al., 2019; Schlissel and Li, 2020) seeks to understand the mechanisms of patterning and cell differentiation through the engineering of genetic circuits (Cachat et al., 2016; Matsuda et al., 2015; Sekine et al., 2018). By re-engineering the Notch/Delta signalling cascade (Figure 1A), Lim and colleagues generated a synthetic circuit capable of reporting and manipulating cell-cell interactions in real time (Morsut et al., 2016). A “sender” cell presenting an extracellular membrane-bound antigen of interest is recognised by a “receiver” cell expressing a chimaeric Synthetic Notch (SynNotch) receptor, composed of an extracellular antigen-recognition domain, the Notch1 core transmembrane domain containing proteolytic cleavage sites, and an intracellular synthetic effector domain (Fig. 1B-D). The modularity of SynNotch circuitry makes it possible to interrogate and manipulate the effects of interactions between cell types of interest.

**Figure 1.**
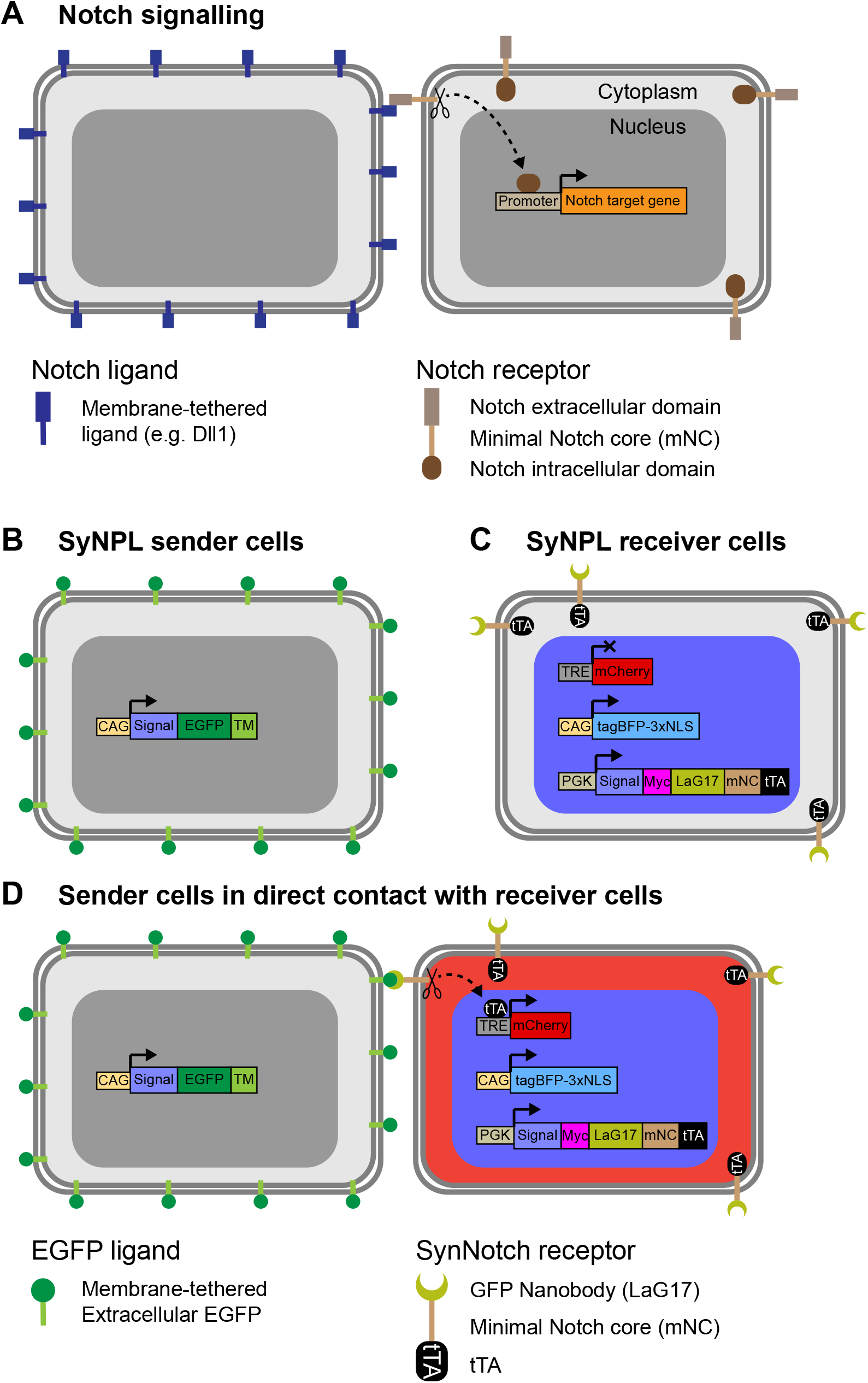
Diagram of SynNotch cell-cell interaction reporter ES cells. (A) Interaction between a membrane-tethered ligand and the Notch receptor extracellular domain leads to the cleavage of the Notch core transmembrane domain, which contains proteolytic cleavage sites. In turn, this leads to the release of the Notch intracellular domain from the membrane, allowing it to translocate to the nucleus and drive transcription of target genes. (B) Sender cells express membrane-tethered extracellular EGFP from a ubiquitous promoter. (C) Receiver cells contain a *TRE-mCherry* transgene, express *tagBFP-3xNLS* from a ubiquitous promoter, and express a SynNotch receptor from a ubiquitous promoter. The SynNotch receptor is comprised of an extracellular LaG17 anti-GFP nanobody, the core transmembrane region of Notch1, and an intracellular tTA. (D) Upon interaction of the EGFP on a sender cell with the SynNotch receptor, the Notch1 core domain is cleaved, releasing the tTA, which can translocate into the nucleus, bind the TRE element, and drive *mCherry* transcription.

SynNotch technology has been used for monitoring cell-cell interactions, generating synthetic patterns, generating synthetic morphogen gradients, inducing contact-mediated gene editing, and generating custom antigen receptor T-cells (Cho et al., 2018; Choe et al., 2021; He et al., 2017; Huang et al., 2020; Roybal et al., 2016; Sgodda et al., 2020; Toda et al., 2018; Toda et al., 2020; Wang et al., 2021). SynNotch technology has been established in *Drosophila* (He et al., 2017) as well as in immortalised cell lines and differentiated cell types, but its potential in the study of mammalian developmental events remains largely untapped.

Mouse embryonic stem cells (ESCs) can be differentiated into any cell type *in vitro*, can give rise to chimaeric embryos and can be used to establish transgenic mouse lines (Bradley et al., 1984; Evans and Kaufman, 1981; Martin, 1981). Adapting the SynNotch system for use in mouse ESCs would therefore permit monitoring and manipulation of cell-cell interactions in a developmental context both *in vivo* and *in vitro*. The original system designed by (Morsut et al., 2016) used lentiviral transduction of immortalised and primary cell lines, where transgene expression was driven from the retroviral SFFV promoter. Lentiviral transduction can lead to multiple copy transgene integration in mouse ESCs (Pfeifer et al., 2002), and the SFFV promoter is prone to silencing in mouse pluripotent cells and their derivatives (Herbst et al., 2012; Pfaff et al., 2013; Wu et al., 2011), making this system suboptimal for mouse ESCs.

In this study, we made several adaptations to the original SynNotch system (Morsut et al., 2016) to establish clonal modular SynNotch pluripotent cell lines (SyNPL). We characterised the SyNPL system by monitoring interactions between EGFP-expressing sender cells and mCherry-inducible receiver cells *in vitro*, then showed that this system can report interactions between neighbouring cells *in vivo* in chimaeric mouse embryos, that it can be used for synthetic patterning, and that its modular design can be exploited to conveniently manipulate cell-cell interactions and drive contact-mediated synthetic cell fate engineering.

## RESULTS

### Design of SyNPL ESCs

We adapted the SynNotch system, which was previously established through viral transduction of immortalised mouse L929 fibroblasts and K562 erythroleukaemic cells (Morsut et al., 2016), for use in mouse ESCs. In this system, sender cells are labelled with membrane-tethered extracellular EGFP (Fig. 1B). Receiver cells constitutively express a SynNotch receptor composed of an anti-GFP nanobody (LaG17) (Fridy et al., 2014), the mouse Notch1 minimal transmembrane core (Uniprot: Q01705, residues 1427-1752), and a tetracycline transactivator (tTA) (Gossen and Bujard, 1992), and contain a tetracycline response element (TRE) promoter capable of driving *mCherry* expression in response to tTA binding (Fig. 1C). Interaction of EGFP on sender cells with the anti-GFP nanobody on receiver cells leads to cleavage of the Notch1 core, releasing the tTA which can translocate to the nucleus, bind to the TRE promoter and drive *mCherry* expression (Fig. 1D). In addition, we constitutively labelled receiver cells with a tagBFP-3xNLS construct (Fig. 1C,D) to conveniently identify them by fluorescence microscopy and flow cytometry even in the absence of a contact-dependent mCherry signal.

Our aims when adapting the SynNotch system were to generate ESC lines with low cell-cell variability, robust and sustained transgene expression, and a modular design to allow convenient transgene exchange. In order to avoid cell-cell variability we generated clonal cell ESC lines with stable genomic integration of the SynNotch system components, delivering transgenes by electroporation rather than lentiviral transduction (Boggs et al., 1986; Charrier et al., 2011; Pfeifer et al., 2002; Smithies et al., 1985). We sought to ensure uniform levels of transgene expression by screening clonal lines and/or targeting transgenes to specific genomic locations, and by replacing the silencing-prone SFFV retroviral promoter used by (Morsut et al., 2016) with CAG (Niwa et al., 1991) or mouse *Pgk1* (McBurney et al., 1991) promoters, which have been extensively characterised in mouse ESCs (Chen et al., 2011; Hong et al., 2007). Finally, we introduced modularity to our system by generating a “landing platform” master cell line to allow recombination-mediated cassette exchange (RMCE) of transgenes of interest.

### Generation of extracellular membrane-tethered EGFP-expressing sender ESCs

We first generated clonal sender cell lines expressing membrane-tethered extracellular EGFP. The CAG and mouse *Pgk1* promoters are both silencing-resistant promoters commonly used to drive ubiquitous transgene expression in ESCs (Herbst et al., 2012; Liew et al., 2007). We asked which of these promoters can generate sender cells with strong and uniform expression of membrane EGFP. We also explored whether EGFP molecules with HA and Myc protein tags can retain “sender” function in pluripotent cells.

We electroporated mouse ESCs with four alternative sender constructs, containing either CAG or mouse *Pgk1* promoters driving expression of either untagged or HA- and Myc-tagged EGFP fused to a membrane-tethering domain (Fig. 2A-D). We isolated and expanded 64 clonal lines derived from stable genomic integration of the four constructs, and screened them by flow cytometry, analysing median EGFP intensity (Fig. 2E,F), percentage of EGFP-positive cells (Fig. 2G,H) and EGFP distribution (Figs. S1, S2). Both CAG and *Pgk1* promoters drive high uniform expression of EGFP, and, as expected, there is considerable variability in EGFP expression between clonal lines.

**Figure 2.**
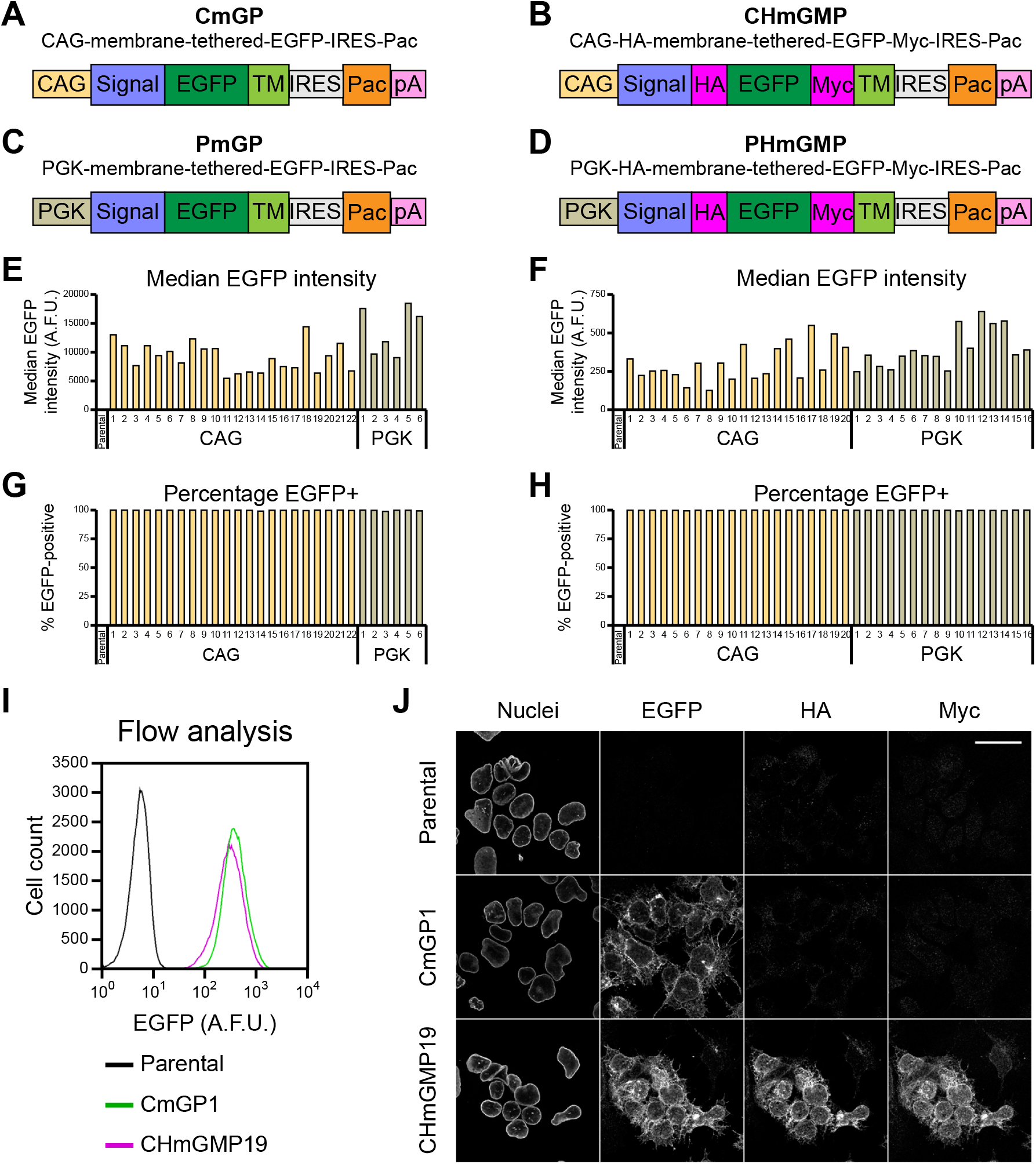
Screening of clonal sender ESC lines. (A-D) Diagram of the constructs used to generate EGFP sender cell lines. (E) Median EGFP intensity and (G) percentage EGFP-positive cells in untagged EGFP sender cells. Parental wild-type cells are included as a negative control. 5000 cells were analysed for each clone. (F) Median EGFP intensity and (H) percentage EGFP-positive cells in HA- and Myc-tagged EGFP sender cells. Parental wild-type cells are included as a negative control. 15000 cells were analysed for each clone. Analyses were performed separately from those in (E), median intensities are not directly comparable. For panels (E-H), n=1. (I) Comparison of EGFP distributions in parental wild-type cells, CmGP clone 1 (CmGP1) and CHmGMP clone 19 (CHmGMP19) sender cells. 85000 cells were analysed for each sample. Data from a single experiment, representative of 9 biological replicates. (J) Immunofluorescence of parental wild-type, CmGP1 and CHmGMP19 sender cells. Scale bar: 30μm. Nuclei: LaminB1. A.F.U.: arbitrary fluorescence units.

We selected one untagged EGFP sender clone (CmGP1) and one HA- and Myc-tagged EGFP sender clone (CHmGMP19), exhibiting high, uniform and similar levels EGFP expression (Fig. 2I) for further analysis. For both clones, the pattern of EGFP expression was consistent with membrane localisation, and, in the case of the HA- and Myc-tagged CHmGMP19 clone, the pattern of HA and Myc expression coincided with that of EGFP (Fig. 2J).

### Generation of a safe harbour site landing pad master ESC line

To facilitate convenient and repeated modification of the genome, we generated a clonal ESC line carrying a “landing pad” targeted to the *Rosa26* locus, a safe harbour site in the mouse genome (Friedrich and Soriano, 1991). This landing pad contains a splice acceptor, the *Neo* (G418/geneticin resistance) gene, and a CAG promoter driving expression of *mKate2-3xNLS*, which encodes a red fluorescent protein with no evident phenotypic effect in mouse embryos (Malaguti et al., 2013; Shcherbo et al., 2009). This entire cassette is flanked by two *attP50* sites, which allows for φC31 integrase-mediated recombination with cassettes flanked by two *attB53* sites (Huang et al., 2009; Tosti et al., 2018) (Fig. S3A). After confirming insertion at the correct genomic locus (Fig. S3B), we verified that all cells express high and uniform levels of mKate2-3xNLS (Fig. S3C,D). We named this cell line EM35.

### An “all-in-one” design fails to generate fully functional mCherry inducible receiver cells

We first asked whether it is possible to target all transcriptional units required for receiver cell activity to the *Rosa26* landing pad in EM35 ESCs, and whether this would lead to the generation of functional receiver ESCs. Design and characterisation of the resulting cell lines is explained in detail in Supplementary Methods and Figs. S4-S9. Briefly, mCherry could, as expected, be induced by subpopulations of tagBFP-positive receiver cells in response to interaction with EGFP-positive sender cells, but this receiver cell design was hampered by variable levels of tagBFP, variable inducibility of mCherry, and low levels of the SynNotch receptor. We conclude that the SynNotch receptor construct and *TRE-mCherry* cassette can function as expected in ESCs, but further modifications to the design are required to obtain a reliable contact-reporting system.

### A multi-step design produces fully functional mCherry-inducible receiver ESCs

We hypothesised that two independent events may be affecting mCherry inducibility in “all-in-one-locus” receiver cells (Figs. S4-S9). Firstly, *mCherry* and *tagBFP* transgenes may have been lost due to mitotic recombination (Stern, 1936) or errors in replication at similar DNA sequences in close proximity (*Pgk1* promoters, *bGHpA* signals). Secondly, the SynNotch receptor may not be expressed at high enough levels (Fig. S9).

We circumvented potential loss of DNA by physically separating the three transcriptional units through random genomic integration of the SynNotch receptor and *tagBFP-3xNLS* cassettes, and by removing identical DNA sequences. To increase levels of SynNotch receptor and obtain uniform levels of tagBFP-3xNLS, we added an internal ribosome entry site (IRES) followed by *Ble* (zeocin resistance gene) downstream of the SynNotch receptor sequence, and an IRES followed by *Hph* (hygromycin B resistance gene) downstream of the *tagBFP-3xNLS* sequence (Fig. 3A).

**Figure 3.**
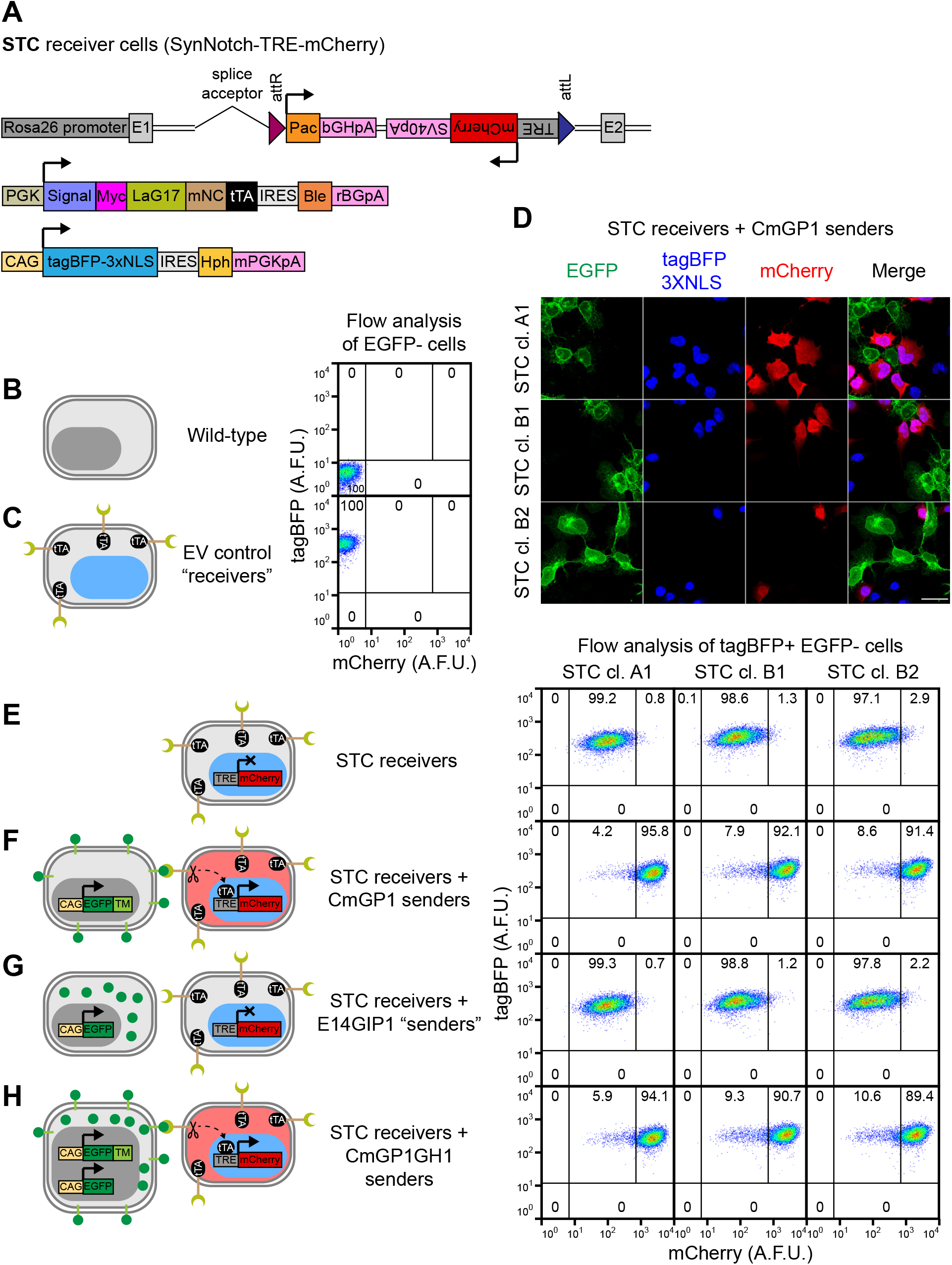
Generation of clonal cell-cell interaction reporter STC receiver cells. (A) Summary of transgenes stably integrated into the genome of STC clonal ESC lines. (B-C) Flow cytometry analysis of mCherry and tagBFP-3xNLS expression in (B) wild-type and (C) control PSNBB-E cells containing all SyNPL receiver constructs except for the TRE-mCherry cassette. These control cell lines were used to set gates for tagBFP- and mCherry-positivity. (D) Immunofluorescence of STC clones A1, B1 and B2 co-cultured with CmGP1 sender cells for 24 hours (1:3 sender:receiver cell ratio). Scale bar: 30μm. (E-H) Flow cytometry analysis of mCherry and tagBFP-3xNLS expression in STC receiver cells cultured alone or in the presence of indicated EGFP-expressing cell lines (9:1 sender:receiver cell ratio). The mCherry-HI gate displayed in figure was set based on mCherry distribution in STC receiver cells cultured alone. Data in these panels were acquired simultaneously to data in panels (B,C) and can be directly compared. For panels (B,C,E-H) percentages of cells in each gate are indicated. 11000 cells were analysed for each sample. Data from a single experiment, representative of 4 biological replicates. Diagrams depicting the transgenes present in each cell type and expected mCherry and tagBFP-3xNLS expression patterns are displayed next to flow cytometry plots.

We first randomly integrated the SynNotch receptor construct into the genome of EM35 landing pad ESCs (Fig. S10A). We selected two clones (35SRZ9, 35SRZ86) with high, uniform Myc expression (Fig. S10B). The levels of Myc in these clones were higher than those in receiver cells generated with an all-in-one design (clones SNCB+4 and SNCB-6) and higher than those in Myc-tagged sender cells (CHmGMP19) (Fig. S10C). We then randomly integrated the *tagBFP-3xNLS* transgene into the genome of these two clones (Fig. S10D). We selected one clone for each parental line with high, uniform expression of tagBFP-3xNLS (PSNB-A clone 10, PSNB-B clone 3) (Fig. S10E,F). We renamed these lines PSNB (<u>P</u>arental <u>S</u>yn<u>N</u>otch tag<u>B</u>FP) clones A and B respectively.

Next, we performed RMCE at the *Rosa26* landing pad in PSNB cells to replace the *mKate2* transgene with one of three constructs: the *TRE-mCherry* cassette present in the SCNB construct, a *tetO-mCherry* cassette with more tTA binding sequences elements in the inducible promoter (to test whether this led to improved mCherry induction), or an empty vector cassette to generate tagBFP-positive mKate2- and mCherry-negative control cell lines (Fig. S11A). We verified that integration of the empty vector cassette led to loss of mKate2 expression, and used these control cell lines to confirm that tagBFP signal was able to unambiguously identify receiver cells (Figs. 3B,C, S11B).

We then asked whether the new receiver cell lines containing inducible mCherry cassettes expressed mCherry in response to co-culture with sender cells. We screened 27 clones for tagBFP and mCherry expression by culturing them in the presence or absence of sender cells for 24 hours (Fig. S11C-G). We observed that all genetically identical clones behaved very similarly, suggesting we were not experiencing silencing or loss of DNA. Clones containing the larger *tetO-mCherry* cassette exhibited high levels of mCherry leakiness in the absence of sender cells. Co-culture with sender cells led to mCherry induction, but the distributions in the presence and absence of sender cells overlapped significantly (Fig. S11D,F,G). Clones containing the smaller *TRE-mCherry* cassette exhibited mCherry leakiness in the absence of sender cells, however co-culture with sender cells led to an increase in mCherry expression to levels which displayed minimal overlap with those seen in cells cultured in the absence of sender cells (Fig. S11C,E,G). Leakiness in the absence of sender cells can be reduced, but not abolished, by treatment of cells with the γ-secretase inhibitor DAPT (which inhibits cleavage of the SynNotch receptor) or with doxycycline (which inhibits tTA-driven transcription) (Fig. S12A,B).

We selected three clones with minimal leakiness and high inducibility for downstream analysis (PSNBA-TRE1, PSNBB-TRE10, PSNBB-TRE9). We renamed these cells STC (for <u>S</u>ynNotch-<u>T</u>RE-m<u>C</u>herry) clones A1, B1 and B2 respectively. These receiver lines induced mCherry after co-culturing sender and receiver cells together for 24 hours. mCherry was specifically induced in tagBFP-positive receiver cells that were in contact with EGFP-positive sender cells (Fig. 3D). We confirmed that mCherry is robustly induced in the majority of tagBFP-positive receiver cells following co-culture with 9-fold excess of CmGP1 sender cells at confluence (Fig 3B,C,E,F).

These observations demonstrate that physical separation of the three transcriptional units in the genome of receiver cells, coupled to the use of internal ribosome entry sites and selectable markers within the units, can lead to the generation of receiver ESC lines which exhibit clear and specific induction of mCherry upon interaction with EGFP-expressing sender cells.

### Extracellular membrane-tethered EGFP is required for contact-mediated transgene induction in receiver cells

It would be useful to make use of existing GFP fluorescent reporter ESCs (for example, cell-state reporters or signalling reporters) to act as sender cells, in order to test how particular cell states may influence direct neighbours. However, many such cell lines make use of non-membrane-tethered GFP, which seems unlikely to interact with the anti-GFP nanobody on STC receiver cells. We therefore wished to test whether membrane tethering of EGFP to the extracellular space was absolutely necessary for effective neighbour-labelling.

We cultured STC receiver cells alone (Fig. 3E), in the presence of CmGP1 sender cells (Fig. 3F), in the presence of a control cell line expressing untagged intracellular EGFP (E14GIP1) (Fig. 3G), or in the presence of CmGP1 sender cells containing an extra untagged intracellular EGFP transgene (CmGP1GH1) (Fig. 3H). E14GIP1 cells, which do not express membrane-tethered EGFP, did not induce mCherry above baseline levels in in co-cultured STC receiver cells (Fig. 3E-G). mCherry is induced to similar levels following co-culture with either CmGP1 or CmGP1GH1 sender cells (Fig. 3F,H), suggesting that the additional untagged EGFP transgene in CmGP1GH1 cells does not interfere with mCherry induction. We conclude that cells containing intracellular GFP cannot function as sender cells unless supplemented with extracellular membrane-tethered EGFP.

### tagBFP-3xNLS lineage label allows identification of EGFP cross-labelled receiver cells

In other cell types, a membrane-tethered anti-GFP nanobody can bind and internalise membrane-tethered GFP on neighbouring cells (Tang et al., 2020). This is also the case in ES cells (Fig. S12B): punctuate EGFP signal is visible in mCherry-expressing activated receiver cells (Fig. S12C). It could therefore be difficult to unambiguously separate of STC receiver cells from CmGP1 sender cells by flow cytometry based on EGFP expression alone (Fig. S12D). This problem is overcome by using the tagBFP-3xNLS lineage label in STC receiver cells (Fig. S12E). Furthermore, separation of sender and receiver cells based on EGFP alone can be achieved by using CmGP1GH1 sender cells, which contain a second EGFP transgene, leading to increased separation between sender cells and cross-labelled receiver cells (Fig. S12F,G).

### Increasing sender:receiver cell ratios leads to increased transgene induction in receiver cells

We next asked how differing sender:receiver cell ratios affect the efficiency of neighbour-labelling. We co-cultured STC receiver cells with different proportions of sender cells for 24 hours (Fig. S13). We observed that as few as 20% of sender cells were sufficient to induce mCherry in approximately half of STC receiver cells, and that 90% sender cells could induce mCherry in over 90% of STC receiver cells (Fig. S13A,B). mCherry fluorescence follows a bimodal distribution in receiver cells exposed to “non-saturating” numbers of sender cells, and a unimodal distribution in receiver cells exposed to “saturating” numbers of sender cells (Fig. S13C-H). This suggests that STC receiver cells which have come into contact with sender cells can uniformly induce high levels of mCherry expression when co-culturing cells at a 9:1 sender:receiver cell ratio (Fig. S13I-K).

### Kinetics of contact-dependent transgene induction in receiver cells

We next performed time-lapse microscopy. In order to capture a range of behaviours, we co-cultured CmGP1GH1 sender cells with STC receiver cells at a 1:1 sender:receiver ratio at moderate density, and filmed cells for 24 hours (Fig. 4A, Movie 1). mCherry first became visible 5-6 hours after initial sender-receiver contact (Fig. 4A, yellow arrowheads). We observed STC receiver cells which did not make contact with sender cells and remained mCherry-negative (Fig. 4A, magenta arrowheads), and an STC receiver cell that made contact with a sender cell 2 hours before the cells were fixed at the 24-hour timepoint for immunofluorescence, and which remained mCherry-negative (Fig. 4A, cyan arrowheads).

**Figure 4.**
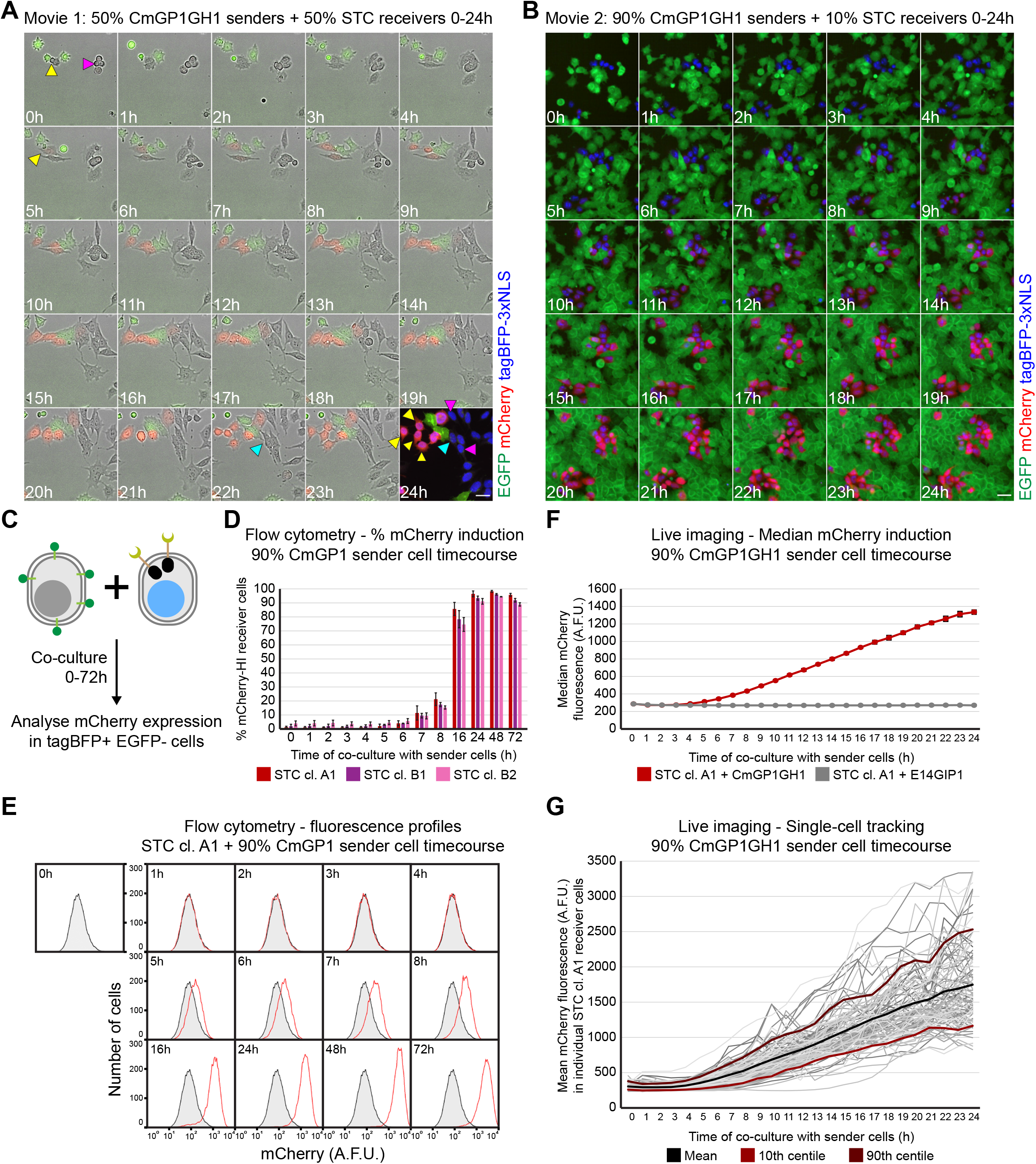
Kinetics of mCherry induction in STC receiver cells. (A) Stills from Movie 1 displaying mCherry and EGFP expression in STC clone A1 receiver cells co-cultured with CmGP1GH1 sender cells (1:1 sender:receiver cell ratio). Immuofluorescence of cells after 24 hours of filming is displayed as a 24 hour timepoint, and includes tagBFP-3xNLS signal in place of a brightfield image. Scale bar: 30μm. Yellow arrowheads: initial STC receiver cell contact with sender cells, onset of mCherry expression, cell descendants at 24 hours. Magenta arrowheads: STC receiver cell not making contact with sender cells, cell descendants at 24 hours. Cyan arrowhead: initial STC receiver cell contact with sender cell, cell descendant at 24 hours. (B) Stills from Movie 2 displaying tagBFP-3xNLS, mCherry and EGFP expression in STC clone A1 receiver cells co-cultured with CmGP1GH1 sender cells (9:1 sender:receiver cell ratio). Scale bar: 30μm. (C) Experimental setup to analyse kinetics of mCherry upregulation in STC receiver cells by flow cytometry. (D) Percentage of mCherry-HI STC receiver cells following co-culture with CmGP1 sender cells for the indicated amount of time (9:1 sender:receiver cell ratio). Data presented as mean ± standard deviation of three independent experiments. A minimum of 8000 cells were analysed for each sample. The mCherry-HI gate was set based on mCherry distribution in STC receiver cells cultured alone. (E) Distribution of mCherry fluorescence in STC clone A1 receiver cells following co-culture with CmGP1 sender cells for the indicated amount of time (9:1 sender:receiver cell ratio). Data from a single experiment, representative of three biological replicates. STC clone A1 cells cultured alone (“0h”) are displayed as a shaded black histogram in all panels. 10000 cells were analysed for each sample. (F) Quantification of live imaging: median mCherry fluorescence intensity in STC clone A1 receiver cells following co-culture with CmGP1GH1 or E14GIP1 cells for the indicated amount of time (9:1 sender:receiver cell ratio). Average of 3 biological replicates, 10 random fields of view/replicate, minimum of 320 cells/replicate/timepoint. Error bars: standard deviation. (G) Mean mCherry fluorescence intensity in individual STC clone A1 receiver cells tracked for 24 hours whilst in co-culture with CmGP1GH1 sender cells (9:1 sender:receiver ratio). Tracks are displayed for 33 randomly selected cells for each of 3 biological replicates (99 cells total). Mean, 10^th^, and 90^th^ centile tracks are also displayed. A.F.U.: arbitrary fluorescence units.

We then quantified the kinetics of mCherry induction. We co-cultured 10% STC receiver cells with 90% sender cells at high density in order to ensure interaction of almost every receiver cell with at least one sender cell (Fig. 4B, Movie 2), and analysed mCherry expression in receiver cells by flow cytometry over the course of 72 hours (Figs. 4C-E, S14A,B), and by live imaging and tracking of individual cells over the course of 24 hours (Figs. 4B,F,G, S14C-H). mCherry is first induced at low levels at around 5 hours (Figs. 4D-G, S14 A-H) and increases until around 48 hours (Figs 4 D-H, S14 A-B).

### Minimum time of contact required for transgene induction

Relying on direct detection of mCherry (Fig 4C-G) is likely to overestimate the minimum duration of cell contact required for mCherry induction because mCherry protein maturation will introduce a time-lag between initiation of mCherry transcription and the detection of mCherry fluorescence. Indeed, time-lapse analysis (Fig. 5A, Movie 3) provides an example of an STC receiver cell which remained in contact with a sender cell for 8 hours, lost contact for 12 hours following a cell division, but continued to increase mCherry expression after losing contact (Fig. 5A, white arrowheads).

**Figure 5.**
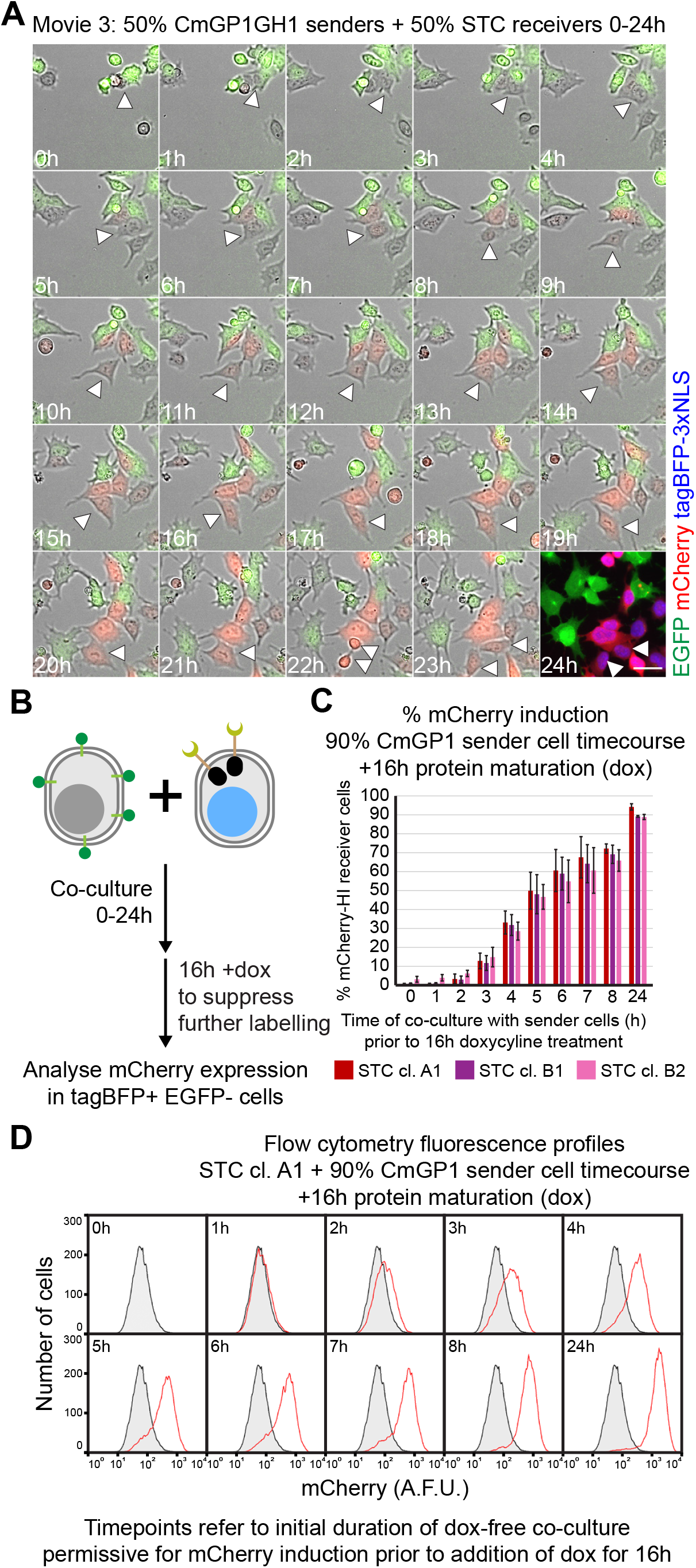
Characterisation of minimal contact time required for mCherry induction in STC receiver cells. (A) Stills from Movie 3 displaying mCherry and EGFP expression in STC clone A1 receiver cells co-cultured with CmGP1GH1 sender cells (1:1 sender:receiver cell ratio). Immuofluorescence of cells after 24 hours of filming is displayed as a 24 hour timepoint, and includes tagBFP-3xNLS signal in place of a brightfield image. Scale bar: 30μm. White arrowheads label an STC receiver cell in contact with sender cells for 7 hours, which then loses contact with sender cells between the 8 and 20 hour timepoints whilst its levels of mCherry keep increasing. (B) Experimental setup to analyse kinetics of mCherry upregulation in STC receiver cells by flow cytometry, allowing time for protein maturation. Following sender:receiver cell co-culture for 0-24 hours, 1μg/ml doxycycline (dox) was added to the culture medium for a further 16 hours in order to inhibit tTA-mediated *mCherry* transcription, and allow translation and folding of previously transcribed *mCherry*. (C) Percentage of mCherry-HI STC receiver cells after co-culture with CmGP1 sender cells for the indicated amount of time and following a further 16 hours of doxycycline treatment (9:1 sender:receiver cell ratio). Data presented as mean ± standard deviation of three independent experiments. A minimum of 8000 cells were analysed for each sample. The mCherry-HI gate was set based on mCherry distribution in STC receiver cells cultured alone in doxycycline for 16 hours. (D) Distribution of mCherry fluorescence in STC clone A1 receiver cells following co-culture with CmGP1 sender cells for the indicated amount of time and 16 hours doxycycline treatment (9:1 sender:receiver cell ratio). Data from a single experiment, representative of three biological replicates. STC clone A1 cells plated with CmGP1 sender cells in doxycycline-containing medium for 16 hours (“0h”) are displayed as a shaded black histogram in all panels. 10000 cells were analysed for each sample.

We designed an experimental strategy to overcome this problem. We co-cultured 10% STC receiver cells with 90% CmGP1 sender cells for various time points between 0 and 24 hours, then added doxycycline to the culture medium for a further 16 hours (Fig. 5B). Doxycycline prevents tTA from binding to *TRE* sequences (Gossen and Bujard, 1992), hence we expect doxycycline administration to halt *mCherry* transcription in receiver cells while still allowing time for mCherry protein to mature. This means that any mCherry signal observed after doxycycline administration should be ascribable to cell-contact-dependent transcription that took place during the initial period of co-culture in doxycycline-free medium. In this experimental setting, we observed low but detectable induction of mCherry when cells had experienced only 2 hours of doxycycline-free co-culture (Figs. 5C,D, S15). These data collectively suggest that 2 hours of sender-receiver contact may be sufficient for induction of low levels of mCherry, and that mCherry levels will keep increasing in receiver cells for a period of time following the loss of sender-receiver contact. This neighbour-labelling system can therefore identify receiver cells that have had relatively brief interactions with sender cells or which have recently lost contact with sender cells.

### Kinetics of contact-dependent mCherry perdurance in receiver cells

We next established how long mCherry signal persists following loss of tTA-mediated *mCherry* transcription. We co-cultured 10% STC receiver cells with 90% CmGP1GH1 cells for 24 hours, then added doxycycline to the culture medium (to block the activity of tTA and halt *mCherry* transcription) and filmed the cells over 48 hours (Fig. S16A,B, Movies 4,5). mCherry fluorescence barely changed for the initial 8-12 hours, then gradually decreased until extinguishment around 38-40 hours after doxycycline addition (Fig. S16A,B). To quantify this process, we co-cultured 10% STC receiver cells with 90% CmGP1 sender cells for 24 hours, then added doxycycline to the culture medium and analysed mCherry fluorescence by flow cytometry at various timepoints (Fig. S16C). No reduction of mCherry signal was observed for the initial 8 hours following doxycycline administration, then median mCherry expression decreased by approximately half at 16 hours, and returned to background levels within 48 hours (Fig. S16D-F). Quantification of live imaging data at hourly timepoints broadly confirmed these observations, with mCherry levels halving after approximately 20-24 hours, and mCherry signal returning to background levels around 48 hours (Fig. S16G).

Taken together, these results suggest that induction of mCherry occurs more rapidly than loss of mCherry signal, presumably due to the high stability of this fluorescent protein, confirming the utility of this system for identifying recent as well as current cell-cell interactions.

### The SyNPL SynNotch cell-cell interaction reporter is functional in early mouse embryos

We asked if the SyNPL system could function *in vivo* in early mouse embryos. We aggregated wild-type morulae with CmGP1GH1 sender cells and/or STC receiver cells, and cultured these to the blastocyst stage (Fig. 6). As expected, all chimaeric blastocysts (80/80) containing both sender and STC receiver cells induced expression of mCherry, whereas no wild-type blastocysts nor blastocysts containing only sender cells displayed mCherry expression (Fig. 6A,B). 18 out of 19 chimaeras containing STC receiver cells alone did not express readily detectable levels of mCherry (Fig. 6B), in line with the low proportion of mCherry-high cells observed *in vitro* in STC receiver cells cultured alone. Treatment of chimaeric embryos with the γ-secretase inhibitor DAPT suppressed mCherry induction, and withdrawal of DAPT allowed mCherry upregulation (Fig. S17), confirming that SynNotch receptor cleavage is required for mCherry induction.

**Figure 6.**
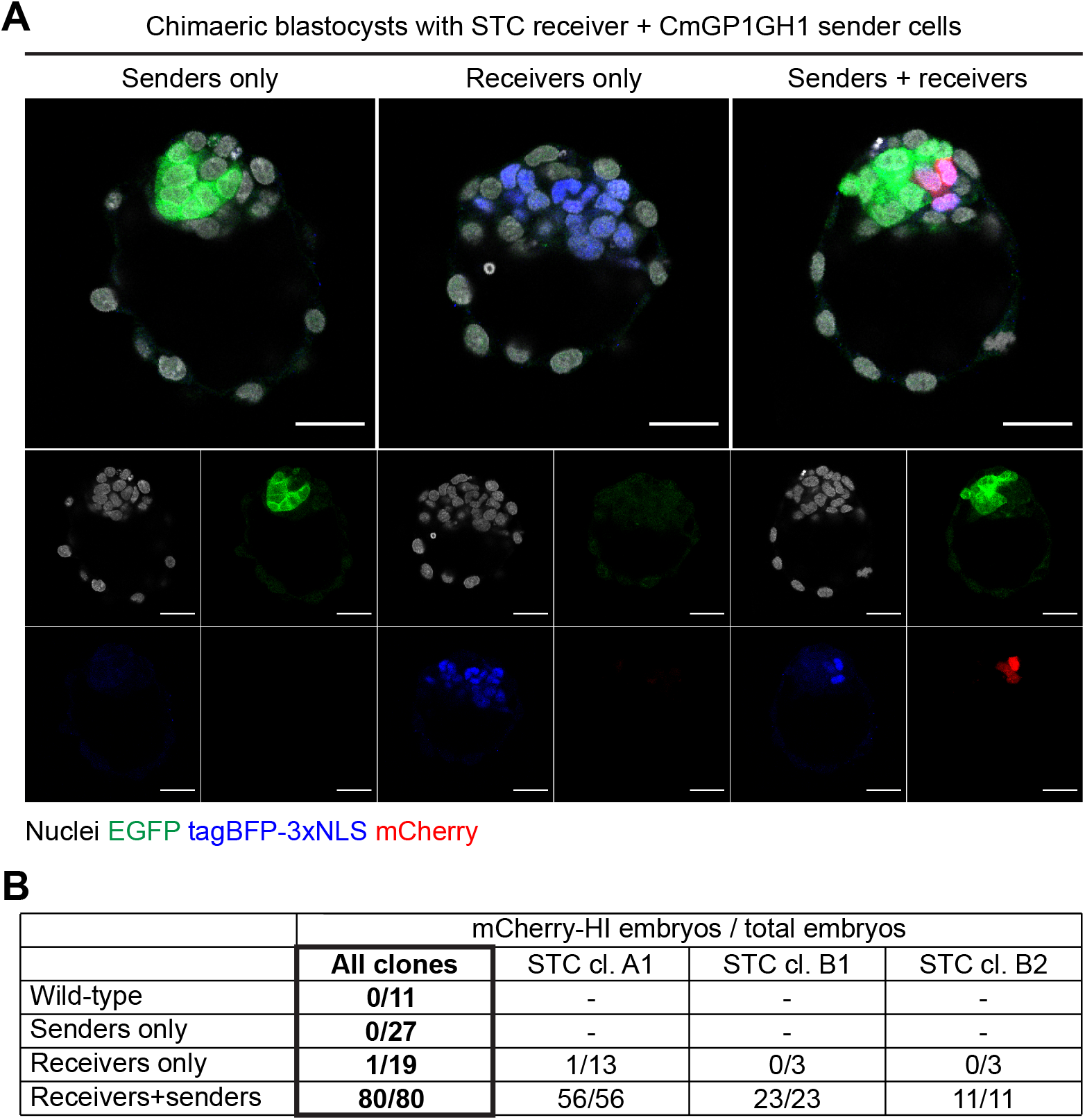
Contact-mediated induction of mCherry in chimaeric blastocysts. (A) Chimaeric blastocysts containing STC clone B1 receiver cells and/or CmGP1GH1 sender cells. The images of the three embryos were taken from the same z-plane of a confocal stack and come from a single field of view. Nuclei were counterstained with DRAQ7. Scale bars: 30μm. (B) Quantification of embryos containing cells expressing readily detectable levels of mCherry (“mCherry-HI”) across all experiments. Embryos containing both sender and receiver cells were used as a reference for scoring sender-only chimaeras, receiver-only chimaeras, and wild-type embryos.

All three STC clonal lines reliably induced mCherry within chimaeric embryos that also contained CmGP1GH1 sender cells (Fig. 5B, Fig. S18A), with mCherry generally appearing within 20h of aggregation (Movie 6). As expected, some receiver cells remain unlabelled when given limited access to sender cells within chimeric blastocysts (aggregations performed with eight receiver cells and only one sender cell: Fig. S18B), in keeping with the contact-dependent nature of SynNotch activation. Post-implantation chimaeras containing both sender and receiver cells displayed mCherry induction throughout the body axis (Fig. S18C). These results suggest that the SyNPL neighbour-labelling system is functional, efficient and reliable *in vivo*.

### Spatial confinement of sender and receiver cells leads to synthetic patterning

SynNotch technology has been successfully employed to generate synthetic patterns. Strategies to achieve this include co-culturing cells in a low sender:receiver cell ratio in order to create 2-dimensional activated receiver cell rings surrounding a clone of sender cells (Morsut et al., 2016), creating self-organising cell aggregates through contact-mediated induction of adhesion molecules (Toda et al., 2018), and recreating morphogen gradients through the use of anchor proteins to capture diffusible receiver cell-activating signal (Toda et al., 2020).

We asked if we could generate a synthetic stripe of transgene expression at the region of contact between sender and receiver cells. We plated CmGP1 sender and STC receiver cells in separate chambers of a removable multi-chamber cell culture insert and allowed them to reach confluence. We then removed the insert, allowing cells to grow in the space between chambers and make contact (Fig. 7A; a detailed description of stripe generation and characterisation can be found in Supplementary Methods and in Fig. S19). A distinct stripe of mCherry expression appeared at the sender:receiver border (Fig. 7B) 24 hours after initial sender:receiver contact. This demonstrates that SynNotch technology can be successfully employed in mouse ESCs to generate synthetic patterns of gene expression.

**Figure 7.**
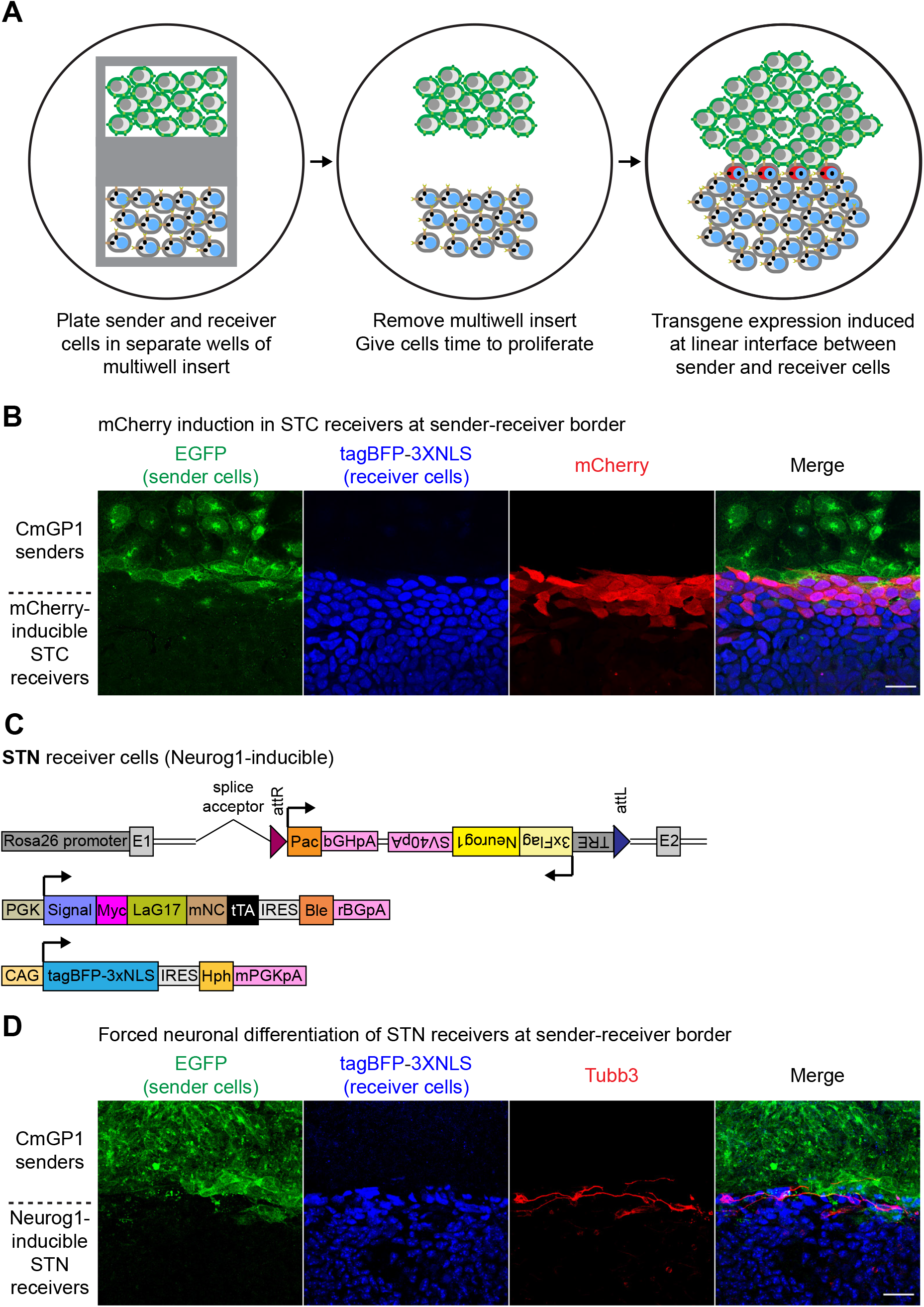
Contact-mediated synthetic patterning of gene expression and fate programming. (A) Diagram illustrating synthetic patterning strategy: sender and receiver cells are grown to confluence in separate chambers of a multi-chamber culture insert. The insert is removed and cells are allowed to proliferate until they come into contact, which induces transgene expression in receiver cells in a stripe pattern. (B) Synthetic striped pattern of mCherry induction in STC clone B1 receiver ESCs in contact with CmGP1 sender ESCs. EGFP, tagBFP, mCherry immunofluorescence. (C) Summary of transgenes stably integrated into the genome of *Neurog1*-inducible STN clonal ESC lines. (D) Synthetic striped pattern of neuronal differentiation of STN receiver ES cells in contact with CmGP1 sender ESCs. EGFP, tagBFP, Tubb3 immunofluorescence. Scale bars: 30μm.

### Harnessing modularity of SyNPL SynNotch ESCs to synthetically alter cell fate

The modularity of our SyNPL SynNotch system design makes it straightforward to generate clonal receiver cell lines with inducible expression of any gene of interest.

The transcription factor Neurog1 (Neurogenin 1) drives neuronal differentiation of progenitor cells during mouse development (Cau et al., 2002; Ma et al., 1998; Yuan and Hassan, 2014). Ectopic expression of Neurog1 is sufficient to drive neuronal differentiation (Cai et al., 2000; Ma et al., 1996) even in mesodermal tissues (Perez et al., 1999) and in mouse ESCs cultured in pluripotent culture conditions (Velkey and O’Shea, 2013). We asked whether a TRE-inducible Neurog1 transgene in receiver cells would drive neuronal differentiation as a specific response to contact with sender cells.

We generated STN (<u>S</u>ynNotch <u>T</u>RE-<u>N</u>eurog1) receiver cells by performing RMCE at the *Rosa26* landing pad in PSNB cell lines to replace the *mKate2-3xNLS* transgene with a *TRE-3xFlag-Neurog1* cassette (Figs. 7C, S20A). We co-cultured STN receiver cells with CmGP1 senders cells for 48 hours, the timepoint at which we observed maximum mCherry induction in STC receiver cells (Figs. 4C,D, S14A,B). We confirmed this resulted in robust induction of 3xFlag-Neurog1 in STN receiver cells compared to STN receiver cells cultured alone (Fig. S20B).

We then asked whether we could induce contact-mediated neuronal differentiation of receiver cells in pluripotent culture conditions, and whether we could engineer differentiation to occur in a synthetic pattern. We repeated the synthetic stripe patterning experiment described above (Fig. 7A,B), using STN receiver cells in place of STC receiver cells. We assessed the expression of the neuronal marker Tubb3 96 hours after initial sender:receiver contact, and observed evident induction of Tubb3 and acquisition of neuronal morphology by STN receiver cells at the sender:receiver border (Fig. 7D). We verified that E14GIP1 cytoplasmic EGFP control cells were unable to induce neuronal differentiation at the border with STN receiver cells (Fig. S21A). We observed that Neurog1 is induced shortly after initial sender:receiver cell contact (Fig. S21B), and that Tubb3 induction first occurs 48 hours after initial interaction between sender and STN receiver cells (Fig. S21C).

We conclude that the interaction between EGFP-expressing sender cells and STN receiver cells can lead to contact-mediated *Neurog1* induction and neuronal differentiation of receiver cells in non-permissive culture conditions. This demonstrates that the SyNPL system can be readily used to generate clonal ESC lines for contact-mediated induction of transgenes of interest, and that these cell lines can in turn be used to manipulate cell-cell interactions in order to program synthetic cell fate decisions in response to contact with a particular cell population at desired locations in space.

## DISCUSSION

Engineering SynNotch machinery (Morsut et al., 2016) into pluripotent cells opens up many opportunities for understanding how direct cell-cell interactions between neighbouring cells can control differentiation decisions, mediate cell competition (Sancho et al., 2013) and orchestrate morphogenesis (Gorfinkiel and Martinez Arias, 2021) as cells differentiate in 2D or 3D culture. Mouse ES cells can contribute to chimaeric embryos, meaning that appropriately engineered cell lines can also be used to understand and control cell-cell interactions during early embryonic development. There are however particular challenges associated with engineering existing SynNotch technologies into pluripotent cells. Here we describe how we overcame these challenges to generate the SyNPL system: a set of clonal SynNotch “sender” and “receiver” mouse ES cells engineered with optimised and modular SynNotch technology. We demonstrate the utility of the SyNPL system for monitoring cell-cell interactions both in culture and in early mouse embryos, and show that we can use this system to engineer contact-dependent cell fate decisions at the boundary between two populations of pluripotent cells.

### Properties of sender cells

We generated sender cell lines expressing high and uniform levels of extracellular membrane-tethered EGFP. This transgenic construct was previously used for SynNotch sender cells (Morsut et al., 2016; Sgodda et al., 2020), and comprises EGFP fused to an N-terminal mouse IgGK signal sequence and a C-terminal human PDGFRB transmembrane domain. The addition of HA and Myc tags at the N- and C-termini of EGFP did not affect the ability of sender cells to induce mCherry expression in STC receiver cells (Fig. S13), so these epitope tags could be helpful for unequivocally identifying and isolating sender cells.

Furthermore, the LaG17 anti-GFP nanobody can also bind to *Aequorea victoria* YFP, CFP and BFP, and *Aequorea macrodactyla* CFP (Fridy et al., 2014), so membrane-tethered versions of these fluorophores could likely be used to induce transgene induction in our receiver cells.

It would be interesting to test whether other cell lines labelled with lipid anchor-tethered GFP (Kondoh et al., 1999; Nowotschin et al., 2009; Rhee et al., 2006; Shioi et al., 2011) could function as SynNotch sender cells; this would require extracellular GFP localisation and generation of sufficient tensile force upon receptor interaction (Morsut et al., 2016). GPI anchors GFP to the outer leaflet of the plasma membrane (Rhee et al., 2006; Sevcsik et al., 2015); GFP-GPI-labelled cells should therefore be capable of acting as sender cells. This does indeed appear to be the case in *Drosophila* (He et al., 2017).

### “All-in-one” locus receiver cells display suboptimal functionality

Attempts at generating “all-in-one”*Rosa26*-targeted receiver cells (termed SNCB+ and SNCB-cells) were unsuccessful. The large variation in tagBFP expression and low proportion of mCherry-inducible ESCs in all clonal lines suggests that either widespread transgene silencing or loss of DNA occurred at the *Rosa26* safe harbour locus in pluripotent cells. Furthermore, rederivation of clonal lines following fluorescence-activated cell sorting of single SNCB+ and SNCB-cells led to re-establishment of the initial heterogeneous distributions of fluorophore expression (Figs. S7, S8), suggesting that the all-in-one design is not optimal for use in pluripotent cells. We were able to overcome these problems by switching to a random-integration strategy, but it is possible that inclusion of IRES-antibiotic resistance cassettes and/or insulator sequences may provide an alternative route towards generating a reliable system without sacrificing the all-in-one-locus approach.

### Landing pad ESCs provide system modularity

Targeting of a landing pad to the *Rosa26* safe harbour locus is an efficient strategy for rapid generation of multiple cell lines through RMCE (Seibler et al., 2005; Tchorz et al., 2012; Tosti et al., 2018). Our cell lines, harbouring a *Rosa26-attP50-Neo-mKate2-attP50* landing pad, make it straightforward to target different transgenes to the same genomic locus in the same parental cell line. Our parental PSNB lines harbouring the *Rosa26* landing pad allowed us to initially test the functionality of SynNotch in ESCs with an inducible *mCherry* transgene in STC receiver cells, prior to generating genetically equivalent STN receiver cells with a *Neurog1* transgene in place of *mCherry*. This modular design therefore makes it possible to readily switch between using SyNPL for monitoring and profiling the consequences of defined cell-cell interactions (based on contact-dependent mCherry expression) and using SyNPL for engineering contact-dependent cell behaviours (based on contact-dependent expression of any cell behaviour-determinant).

### Describing the properties of the SyNPL system

We characterised various aspects of the SyNPL system that will help inform the experimental design for users of these cells.

Previous studies have co-cultured senders and receiver cells at different ratios (ranging from 1:50 to 5:1), and for varying times (ranging from 10 minutes to several days) (Cho et al., 2018; Choe et al., 2021; He et al., 2017; Huang et al., 2020; Luo et al., 2019; Matsunaga et al., 2020; Morsut et al., 2016; Roybal et al., 2016; Sgodda et al., 2020; Srivastava et al., 2019; Toda et al., 2018; Toda et al., 2020; Wang et al., 2021; Yang et al., 2020). We analysed transgene induction in STC receiver cells at 11 different sender:receiver cell ratios (ranging from 1:19 to 9:1), and observed that higher proportions of sender cells in culture result in a higher proportion of receiver cells inducing mCherry. This is in line with the results obtained by (Sgodda et al., 2020) when comparing three different sender:receiver cell ratios, and with the observations of (Morsut et al., 2016), who exposed receiver cells to varying concentrations of sender ligand. By finely varying the concentrations of sender ligand, Lim and colleagues described the transgene induction response as sigmoidal (Morsut et al., 2016), which was not evident in our data. It is however possible that by testing lower sender:receiver cell ratios this might hold true in our SynNotch system too.

When analysing transgene expression within single samples, we found that mCherry distribution follows a bimodal on/off response, indicative of the presence of receiver cells which do not interact with sender cells at low sender:receiver cell ratios. This bimodal pattern of transgene induction is also evident in data from Cantz and colleagues (Sgodda et al., 2020). The ability of individual sender cells to induce mCherry induction in STC receiver cells (as seen in Movies 1 and 3) implies that this system can be effectively employed to study the effect of interactions between receiver cells and individual and/or rare sender cells in relevant model systems.

We performed a high-resolution study of the kinetics of mCherry induction and downregulation in STC receiver cells. We observed low levels of mCherry induction in STC receiver cells following 2 hours of co-culture with sender cells, provided we allowed time for subsequent protein maturation, and observed maximum mCherry induction following 48 hours of co-culture. Previous studies making use of lentiviral-delivered transgenes suggest that 10 minutes may be sufficient for transgene activation in HEK293 receiver cells, and that 1 hour may be sufficient for transgene induction in L929 receiver cells, as long as protein maturation time is allowed (Morsut et al., 2016; Sgodda et al., 2020). This is significantly faster than what we observed in this study, and may be ascribable to lentiviral transduction leading to higher levels of SynNotch receptor and/or integration of multiple transgene copies compared to our clonal mouse ESC lines.

In our system, mCherry downregulation did not commence for at least 8 hours following simulated loss of sender:receiver cell contact, with full loss of signal occurring after more than 40 hours. This is in line with previous observations in L929 receiver cells, where inducible GFP transgene expression was lost between 24 and 50 hours after sender cells were removed from culture (Morsut et al., 2016). The kinetics of mCherry induction and downregulation suggest that this SynNotch system is suited for the study of cell-cell interactions with a temporal range of hours rather than minutes, and that “memory” of such interactions will persist for a few days. mCherry signal intensity will be influenced not only by the duration of contact but also, where cells have moved apart, by the time elapsed since last contact: this may complicate interpretation of data from this system for some applications. Should this persistence of mCherry signal prove inconvenient for the study of particular processes, the PSNB landing pad parental cell lines can be used to readily generate cell interaction reporter receiver cells harbouring destabilised inducible transgenes with short half-lives.

### Exploring the roles of cell-cell interactions *in vivo* and *in vitro*

We demonstrated that our clonal mouse ESC lines can be used *in vivo* in chimaeric embryos. The ability to conveniently switch between *in vitro* and *in vivo* experimentation was a key reason for us to establish SynNotch technology in mouse ESCs. Both the receiver lines we generated and the parental PSNB landing pad cell lines offer the power and flexibility to address questions we have so far been unable to answer in *in vitro* and *in vivo* settings. For example, the system could be employed in cell competition studies: STC receiver cells could be used to identify and isolate the direct neighbours of EGFP-tagged “loser” cells, and profiled to study what changes are induced upon interaction with loser cells in order to bring about their elimination. Receiver cells could also be engineered to express candidate fitness-altering transgenes in response to interaction with EGFP-tagged wild-type sender cells, as successfully demonstrated in *Drosophila* by (He et al., 2017).

The establishment of this system in mouse ESCs also allows to monitor and manipulate the effects of cell-cell interactions in specific cell types obtained through directed differentiation. We also demonstrated that our cell lines can be used to generate synthetic patterns of gene expression, resulting in spatially-defined programming of cell fate. The combination of directed differentiation of ESCs, spatial confinement of sender and receiver cells, and contact-mediated cell fate engineering provides many possibilities for the study of cell-cell interactions in any developmental process of interest.

### Concluding remarks

Cell-cell interactions are a shared feature of the development of all multicellular organisms. Whilst the particulars of these interactions vary greatly amongst eukaryotic supergroups, it is clear that they play an essential role in development (Armingol et al., 2021). The Synthetic Biology field has recently developed several applications to monitor cell communication, such as SynNotch (Morsut et al., 2016) and derivative systems (Zhu et al., 2021), direct transfer of fluorophores to neighbouring cells (Ombrato et al., 2019; Tang et al., 2020), reconstitution of a fluorophore following interaction between different cell types carrying non-fluorescent fluorophore fragments (Kinoshita et al., 2020). We have here demonstrated how SynNotch technology can be used to monitor and manipulate cell-cell interactions in mouse ESCs and in mouse embryos.

## MATERIALS AND METHODS

### Reagents

| REAGENT | SOURCE | IDENTIFIER | CONCENTRATION |
| --- | --- | --- | --- |
| Antibodies and nuclear counterstain |  |  |  |
| Mouse monoclonal anti-Flag | Sigma-Aldrich | Cat# F3165;<br>RRID:<br>AB_259529 | 1:1000 |
| Chicken polyclonal anti-GFP | Abcam | Cat# ab13970;<br>RRID:<br>AB_300798 | 1:1000 |
| Goat polyclonal anti-Myc | Abcam | Cat# ab9132;<br>RRID:<br>AB_307033 | 1:1000 |
| Mouse monoclonal anti-Myc<br>AlexaFluor 488 conjugate | Santa Cruz | Cat# sc-40<br>AF488; RRID:<br>AB_2892598 | 1:200 |
| Rabbit polyclonal anti-LaminB1 | Abcam | Cat#ab16048;<br>RRID:<br>AB_443298 | 1:1000 |
| Rat monoclonal anti-mCherry | Invitrogen | Cat# M11217;<br>RRID:<br>AB_2536611 | 1:1000 |
| Mouse monoclonal anti-Nup107 (NPC) | Abcam | Cat# ab24609;<br>RRID:<br>AB_448181 | 1:1000 |
| Rabbit polyclonal anti-tRFP (tagBFP) | Evrogen | Cat# AB233;<br>RRID:<br>AB_2571743 | 1:1000 |
| Mouse monoclonal anti-Tubb3 | Biolegend | Cat# 801201;<br>RRID:<br>AB_2313773 | 1:1000 |
| DRAQ7 | Abcam | Cat# ab109202 | 1µM (IF); 300nM<br>(flow cytometry) |
| DAPI | Biotium | Cat# 40043 | 1µM |
| Chemicals and solutions |  |  |  |
| 2-Mercaptoethanol | Gibco | Cat# 31350010 | 100nM |
| Accutase | Sigma-Aldrich | A6964 |  |
| Benzyl alcohol | Alfa Aesar | Cat# 100-51-6 |  |
| Benzyl benzoate | Sigma | Cat# B9550 |  |
| Chorulon (hCG) | Intervet | Cat# CH-475-1 | 100 IU/ml |
| DAPT | Abcam | Cat# ab120633 | 100µM |
| DMEM, high glucose, no glutamine, no<br>phenol red | Gibco | Cat# 31053028 |  |
| Donkey serum | Sigma-Aldrich | Cat# D9663 | 3% v/v |
| Doxycycline hyclate | Sigma-Aldrich | Cat# D9891 | 1µg/ml |
| Dulbecco's phosphate buffered saline<br>(PBS) | Sigma-Aldrich | Cat# D8537 |  |
| Fibronectin from bovine plasma<br>solution | Sigma-Aldrich | Cat# F1141 |  |
| Foetal calf serum | Life Technologies | Cat# 10270106 | 10% v/v |
| Formaldehyde 37-41% | Fisher Scientific | Cat# F/1501/PB08 | 4% w/v |
| Gelatin | Sigma-Aldrich | Cat# G1890 | 0.1% w/v |
| Geneticin (G418) | Gibco | Cat# 11811031 | 200µg/ml |
| Glasgow Minimum Essential Medium | Sigma-Aldrich | Cat# G5154 |  |
| Hygromycin B | Gibco | Cat# 10687010 | 200µg/ml |
| L-Glutamine | Thermo Scientific | Cat# 25030024 | 2mM |
| Lipofectamine 3000 transfection reagent | Life Technologies | Cat# L3000008 |  |
| M2 medium | Sigma-Aldrich | Cat# M7167 |  |
| MEM Non-essential amino acids solution (100X) | Gibco | Cat# 11140050 | 1X |
| Penicillin/streptomycin | Gibco | Cat# 15140122 | 100 U/ml |
| PMSG | ProSpec | Cat# HOR-272 | 100 IU/ml |
| Prolong Gold Antifade Mountant | Thermo Fisher | Cat# P36930 |  |
| Puromycin dihydrochloride | Sigma-Aldrich | Cat# P8833 | 2µg/ml |
| Sodium pyruvate | Gibco | Cat# 11360039 | 1 mM |
| Triton X-100 | Sigma-Aldrich | Cat# X100 | 0.1% v/v |
| Trypsin EDTA 0.25% | Gibco | Cat# 25200072 | 0.05% w/v |
| Tyrode's solution, acidic | Sigma-Aldrich | Cat# T1788 |  |
| Zeocin | Gibco | Cat# R25001 | 100µg/ml |
| Cell culture substrates and inserts |  |  |  |
| 3 well silicone culture insert | Ibidi | Cat# 80369 |  |
| µ-Slide 8 Well | Ibidi | Cat# 80826 |  |
| µ-Slide 4 Well | Ibidi | Cat# 80426 |  |
| µ-Plate 96 Well Black | Ibidi | Cat# 89626 |  |
| µ-Slide Angiogenesis, uncoated | Ibidi | Cat# 81501 |  |
| Glass coverslip, 24mm diameter | VWR | Cat# 631-1583 |  |
| Plastic flasks and plates for routine cell culture | Corning | Various |  |
| DNA constructs |  |  |  |
| pHR_EGFPligand | (Morsut et al., 2016) | Addgene 79129 |  |
| pHR_SFFV_LaG17_synNotch_TetRVP64 | (Morsut et al., 2016) | Addgene 79128 |  |
| pDisplay-EGFP-TM | (Han et al., 2004) |  |  |
| pHR_TRE-mCherry-PGK-tagBFP-WPRE | Elise Cachat, University of Edinburgh |  |  |
| CAG-φC31 integrase | (Monetti et al., 2011) | Similar plasmid available: Addgene 62658 |  |
| pRosa26-DEST-1lox | (Tosti et al., 2018) |  |  |
| pENTR-2xAttP50 | (Tosti et al., 2018) |  |  |
| pENTR-2xAttB53 | (Tosti et al., 2018) |  |  |
| pPyCAG-EGFP-TM-IRES-Pac | This study | Addgene 183603 |  |
| pPyPGK-EGFP-TM-IRES-Pac | This study | Addgene 183604 |  |
| pPyCAG-HA-EGFP-Myc-TM-IRES-Pac | This study | Addgene 183605 |  |
| pPyPGK-HA-EGFP-Myc-TM-IRES-Pac | This study | Addgene 183606 |  |
| pPyPGK-Myc-LaG17-Notch1TM-tTA-IRES-Ble | This study | Addgene 183607 |  |
| pPyCAG-tagBFP-3xNLS-IRES-Hph | This study | Addgene 183608 |  |
| pmROSA26-attP50-Neo-mKate2-3xNLS-attP50 | This study | Addgene 183609 |  |
| attB53-Pac-attB53 | This study | Addgene 183610 |  |
| attB53-Pac-TRE-mCherry-attB53 | This study | Addgene 183611 |  |
| attB53-Pac-tetO-mCherry-attB53 | This study | Addgene 183612 |  |
| attB53-Pac-TRE-3xFlag-Neurog1-attB53 | This study | Addgene 183613 |  |
| attB53_SNCB+_attB53 | This study | Addgene 183614 |  |
| attB53_SNCB-_attB53 | This study | Addgene 183615 |  |
| Cell lines |  |  |  |
| E14Ju09 mouse ESCs (129/Ola, male) | (Hamilton and Brickman, 2014) |  |  |
| CmGP1 sender mouse ESCs (129/Ola, male) | This study |  |  |
| CHmGMP19 sender mouse ESCs (129/Ola, male) | This study |  |  |
| CmGP1GH1 sender mouse ESCs (129/Ola, male) | This study |  |  |
| E14GIP1 mouse ESCs (129/Ola, male) | This study |  |  |
| EM35 mouse ESCs (129/Ola, male) | This study |  |  |
| SNCB+ receiver mouse ESCs (129/Ola, male) | This study |  |  |
| SNCB- receiver mouse ESCs (129/Ola, male) | This study |  |  |
| 35SRZ mouse ESCs (129/Ola, male) | This study |  |  |
| PSNB mouse ESCs (129/Ola, male) |  |  |  |
| PSNB-E mouse ESCs (129/Ola, male) | This study |  |  |
| PSNB-tetO receiver mouse ESCs (129/Ola, male) | This study |  |  |
| STC receiver mouse ESCs (129/Ola, male) | This study |  |  |
| STN receiver mouse ESCs (129/Ola, male) | This study |  |  |

### Animal care and use

Animal experiments were performed under the UK Home Office project license PEEC9E359, approved by the Animal Welfare and Ethical Review Panel of the University of Edinburgh and within the conditions of the Animals (Scientific Procedures) Act 1986.

### Chimaera generation

C57BL/6 female mice (Charles River) were superovulated (100 IU/ml PMSG and 100 IU/ml hCG intraperitoneal injections 48 hours apart) and crossed with wild-type stud male mice. Pregnant mice were culled at embryonic day 2.5 (E2.5) by cervical dislocation, ovaries with oviducts were dissected and collected in pre-warmed M2 medium. Oviducts were flushed using PBS and a 20-gauge needle attached to a 1ml syringe and filled with PB1 (Whittingham, 1974). E2.5 embryos were collected and washed in PB1, their zona pellucida was removed using acidic Tyrode’s solution, and they were transferred to a plate with incisions where two clumps of approximately 8 sender and 8 receiver cells were added to each embryo. Embryos were then incubated at 37°C in 5% CO_2_ for 48 hours prior to fixation, or for 24 hours prior to transfer to pseudopregnant females for the generation of post-implantation chimaeras. For DAPT treatment experiments, DAPT was equilibrated for several hours at 37°C prior to addition to embryos in order to avoid precipitation. Embryos subject to DAPT withdrawal were washed twice before being placed in DAPT-free medium. The sex of embryos used in this study was not determined.

### Mouse ESC culture

Mouse embryonic stem cells were routinely maintained on gelatinised culture vessels (Corning) at 37°C and 5% CO_2_ in Glasgow Minimum Essential Medium (GMEM) supplemented with 10% foetal calf serum (FCS), 100U/ml LIF (produced in-house), 100nM 2-mercaptoethanol, 1X non-essential amino acids, 2mM L-Glutamine), 1mM Sodium Pyruvate (medium referred to as “ES cell culture medium” or “LIF+FCS”). The medium was supplemented with 200μg/ml G418, 2μg/ml puromycin, 200μg/ml hygromycin B and/or 100μg/ml zeocin as appropriate.

For live imaging, GMEM was replaced with phenol red-free Dulbecco’s Modified Eagle Medium (DMEM), with all other components of the culture medium used at identical concentrations.

### DNA constructs

pHR_SFFV_LaG17_synNotch_TetRVP64 (Addgene plasmid #79128) (Morsut et al., 2016) and pHR_EGFPligand (Addgene plasmid #79129) (Morsut et al., 2016) were kind gifts from Dr Wendell Lim (UCSF) and Dr Leonardo Morsut (USC). pDisplay-GFP-TM (Han et al., 2004) was a kind gift of Dr Luis Ángel Fernández (CNB-CSIC). CAG-ϕC31 integrase (Monetti et al., 2011) was a kind gift of Dr Andras Nagy (Lunenfeld-Tanenbaum Research Institute). pHR_TRE-mCherry-PGK-tagBFP-WPRE was a kind gift of Dr Elise Cachat (The University of Edinburgh). pRosa26-DEST-1lox, pENTR-2xAttP50 and pENTR-2xAttB53 (Tosti et al., 2018) constructs were kind gifts of Dr Keisuke Kaji (The University of Edinburgh).

Untagged transmembrane EGFP constructs were generated by digesting pHR_EGFPligand with XhoI+NotI, and ligating the *IgGK signal-EGFP-PDGFRB TMD* cassette into XhoI+NotI-digested *pPyCAG-IRES-Pac* (Malaguti et al., 2019) or *pPyPGK-IRES-Pac* (Rao et al., 2020) vector backbones. HA- and Myc-tagged EGFP constructs were generated by PCR-amplifying an *IgGK signal-HA-EGFP-Myc-PDGFRB TMD* cassette flanked by PspXI and NotI sites from pDisplay-GFP-TM, digesting the amplicon with PspXI+NotI, and ligating the insert into XhoI+NotI-digested *pPyCAG-IRES-Pac* or *pPyPGK-IRES-Pac* vector backbones.

The *pPyPGK-CD8a signal-Myc-LaG17-Notch1 minimal transmembrane core-tTA-IRES-Ble* SynNotch receptor construct was generated by PCR-amplifying a *CD8a signal-Myc-LaG17-Notch1 minimal transmembrane core-tTA* cassette flanked by XhoI and Bsu36I sites from pHR_SFFV_LaG17_synNotch_TetRVP64, digesting the amplicon with XhoI+Bsu36I, and ligating the insert into a XhoI+Bsu36I-digested *pPyPGK-IRES-Ble* vector backbone. The mouse Notch1 minimal transmembrane core consists of residues 1427-1752 (Uniprot: Q01705).

The *pPyCAG-tagBFP-3xNLS-IRES-Hph* construct was generated by PCR-amplifying a *tagBFP* cassette flanked by XhoI and KasI sites from pHR_TRE-mCherry-PGK-tagBFP-WPRE, digesting the amplicon with XhoI+KasI, and ligating the insert into a XhoI+NotI-digested *pPyCAG-IRES-Hph* backbone (Malaguti et al., 2019) alongside oligonucleotides annealed to generate a *3xNLS* fragment with KasI and NotI overhangs (Malaguti et al., 2013).

The *Rosa26* landing pad targeting vector was generated by Gateway Cloning (Invitrogen) of an *attL1-attP50-Neo-SV40pA-(CAG-mKate2-3xNLS-bGHpA)-attP50-attL2* cassette into the pRosa26-DEST-1lox targeting vector. Its final structure is as follows: *Rosa26 5’HA-splice acceptor-loxP-attP50-Neo-SV40pA-(CAG-mKate2-3xNLS-bGHpA)-attP50-Rosa26 3’HA-PGK-DTA-bGHpA.*Sequence in brackets is on -strand.

An *attB53-Pac-attB53*“empty vector” construct for RMCE at the *Rosa26* locus was generated by adding a *Pac-bGHpA* cassette followed by an EcoRV restriction site to pENTR-2xAttB53 by Gibson assembly.

The attB53-TRE-mCherry-attB53 RMCE construct used to generated STC receiver cells from PSNB landing pad lines was generated by PCR-amplifying a *TRE-mCherry-SV40pA* cassette flanked by EcoRV-AscI and EcoRV-BamHI sites from pHR_TRE-mCherry-PGK-tagBFP-WPRE, digesting the amplicon with EcoRV, ligating the insert into EcoRV-digested *attB53-Pac-attB53* backbone, and screening for insertion on the -strand. The attB53-tetO-mCherry-attB53 RMCE construct used to generate PSNB-tetO cells from PNSB landing pad lines was generated by PCR-amplifying a *tetO-mCherry-rBGpA* cassette flanked by MluI and BamHI sites, digesting the amplicon with MluI+BamHI and ligating the insert into AscI+BamHI-digested *attB53-Pac-TRE-mCherry-attB53*.

The attB53-TRE-3xFlag-Neurog1-attB53 RMCE construct used to generate STN receiver cells from PSNB landing pad lines was generated by PCR-amplifying a *3xFlag-Neurog1* cassette flanked by NdeI and MfeI sites from wild-type mouse cDNA, digesting the amplicon with NdeI+MfeI and ligating the insert into NdeI+MfeI-digested *attB53-Pac-TRE-mCherry-attB53* backbone (in which *mCherry* is flanked by NfeI and MfeI sites).

attB53_SNCB+_attB53 and attB53_SNCB_attB53 constructs were generated in two steps. First, the base *attB53-Pac-bGHpA-attB53* RMCE construct was linearised with EcoRV, and ligated with a HincII-*PGK-CD8a signal-Myc-LaG17-Notch1 minimal transmembrane core-tTA-*HindIII fragment (digested from the SynNotch receptor construct described above) and a HindIII-*bGHpA*-EcoRV fragment, and clones were screened for insertion of the SynNotch receptor on the +strand. Next, the resulting construct was linearised with EcoRV, and ligated with an EcoRV-*TRE-mCherry-SV40pA-PGK-tagBFP*-PacI fragment (digested from pHR_TRE-mCherry-PGK-tagBFP-WPRE) and a PacI-*bGHpA*-EcoRV fragment. Correct assembly on the + and - strands generated the attB53_SNCB+_attB53and attB53_SNCB-_attB53 constructs respectively.

**Abbreviations**
TMD: transmembrane domain
IRES: internal ribosome entry site
5’/3’ HA: 5’/3’ homology arm
SV40pA: SV40 polyadenylation signal sequence
bGHpA: bovine growth hormone polyadenylation signal sequence
rBGpA: rabbit beta-globin polyadenylation signal sequence
NLS: nuclear localisation signal
PGK: mouse *Pgk1* promoter
Pac: puromycin N-acetyltransferase
Hph: hygromycin B phosphotransferase
Ble: bleomycin resistance gene (confers zeocin resistance)
tTA: tetracycline transactivator

Plasmids that we generated for this study will be made available on Addgene.

### Transfections

For electroporations, 10^7^ ESCs were electroporated with 100μg DNA using a BioRad GenePulser set to 800V/3μF. For nucleofections, 5×10^5^ ESCs were nucleofected with 5μg DNA with the Lonza P3 Primary Cell Nucleofector Unit and kit, using program CG-104, and following manufacturer instructions. For lipofections, 10^5^ ESCs were lipofected with 3μg DNA mixed with 3μl Lipofectamine 3000 and 6μl P3000 solution, following manufacturer instructions. For ϕC31-mediated RMCE, equal masses of RMCE constructs and CAG-ϕC31 vector were transfected.

Clonal ESC lines were generated by transfecting constructs of interest into ESCs, then plating cells at low density onto gelatinised 9cm dishes in the absence of selection. Selective medium was added 48 hours post-transfection and replaced every other day. After 7-10 days, clones were manually picked, dissociated, and replated into gelatinised 96-well culture plates. Clones were transferred to gelatinised vessels with larger culture areas when confluent, screened as appropriate, expanded and cryopreserved.

### Cell lines

E14Ju09 ESCs are a 129/Ola male wild-type clonal line derived from E14tg2a (Hamilton and Brickman, 2014; Hooper et al., 1987).

Sender cells were generated by electroporating E14Ju09 ESCs with one of four constructs: *pPyCAG-IgGKsignal-EGFP-PDGFRB TMD-IRES-Pac* (CmGP cells), *pPyPGK-IgGK signal-EGFP-PDGFRB TMD-IRES-Pac* (PmGP cells), *pPyCAG-IgGK signal-HA-EGFP-Myc-PDGFRB TMD-IRES-Pac* (CHmGMP cells), *pPyPGK-IgGK signal-HA-EGFP-Myc-PDGFRB TMD-IRES-Pac* (PHmGMP cells). Simplified versions of the constructs are displayed in Fig. 2. HA and Myc tags were used in CHmGMP and PHmGMP cell lines as additional markers to identify sender cells, but are not essential given that sender cells can be identified by GFP fluorescence. CmGP1GH1 sender cells were generated by lipofecting CmGP1 sender cells with a *pPyCAG-EGFP-IRES-Hph* construct. E14GIP1 “cytoplasmic sender” cells were generated by lipofecting E14Ju09 ESCs with a *pPyCAG-EGFP-IRES-Pac* construct.

EM35 landing pad cells were generated by electroporating E14Ju09 ESCs with the *Rosa26* landing pad targeting vector described above. Correct targeting was verified by genomic DNA PCR with the following primers: Forward: GGCGGACTGGCGGGACTA, Reverse:

GGGACAGGATAAGTATGACATCATCAAGG. Primer locations and expected band sizes are displayed in Fig. S3A. This PCR strategy was modified from that described by (Mort et al., 2014) to suit the different sequence of our *Rosa26* targeting vector.

SNCB+ and SNCB-receiver cells were generated by electroporating EM35 ESCs with the constructs depicted in Figs. S4A, S6A, and a *CAG-φC31 integrase* construct to mediate RMCE.

35SRZ landing pad cells were generated by electroporating EM35 landing pad cells with *pPyPGK-CD8a signal-Myc-LaG17-Notch1 minimal transmembrane core-tTA-IRES-Ble.*

PSNB landing pad cell lines were generated by nucleofecting 35SRZ ESCs with a *pPyCAG-tagBFP-3xNLS-IRES-Hph* construct. PSNB-A cells were derived from 35SRZ clone 9 (PSNB-A clone 10 renamed PSNB clone A), PSNB-B cells were derived from 35SRZ clone 86 (PNSB-B clone 3 renamed PSNB clone B).

STC receiver cells were generated by nucleofecting PSNB ESCs with *CAG-φC31 integrase* and the following RMCE construct: *attB53-Pac-bGHpA-(TRE-mCherry-SV40pA)-attB53.* Sequence in brackets is on -strand. STC clone A1 was derived from PSNB clone A, STC clones B1 and B2 were derived from PSNB clone B.

PSNB-tetO cells were generated by nucleofecting PSNB ESCs with *CAG-φC31 integrase* and the following RMCE construct: *attB53-Pac-bGHpA-(tetO-mCherry-SV40pA)-attB53.* Sequence in brackets is on -strand.

PNSB-E cells were generated by nucleofecting PSNB ESCs with *CAG-φC31 integrase* and the following RMCE construct: *attB53-Pac-bGHpA-attB53*.

STN receiver cells were generated by nucleofecting PSNB clone A ESCs with *CAG-φC31 integrase* and the following RMCE construct: *attB53-Pac-bGHpA-(TRE-3xFlag-Neurog1-SV40pA)-attB53.* Sequence in brackets is on -strand.

Cell lines generated in this study were routinely karyotyped by chromosome count, checked for absence of mycoplasma infection, and are available on request.

### Co-culture experiments

Sender and receiver cells were detached from culture vessels with accutase, quenched in ESC culture medium, pelleted by spinning at 300g for 3 minutes, resuspended in ESC culture medium supplemented with 2μg/ml puromycin and counted. Cells were plated at ratios described in figure legends, and at empirically determined optimal densities.

For flow cytometry experiments, cells were plated onto 12-well plates coated with 7.5μg/ml fibronectin, at the following densities: experiments carried out in the absence of doxycyline: 1h-8h: 4×10^5^ cells/well; 16h: 2.4×10^5^ cells/well; 24h: 1.6×10^5^ cells/well; 48h: 8×10^4^ cells/well; 72h: 4×10^4^ cells/well. mCherry induction with 16h doxycycline (Fig. 4F-H): 0-4h: 2.4×10^5^ cells/well; 5-8h: 1.6×10^5^ cells/well; 24h: 8×10^4^ cells/well. mCherry downregulation experiments (Fig. S14): 0h-8h: 1.6×10^5^ cells/well; 16h: 1.2×10^5^ cells/well; 24h: 8×10^4^ cells/well; 48h: 4×10^4^ cells/well.

For immunofluorescence experiments, cells were plated on flamed 24mm glass coverslips housed in a 6-well plate coated with 7.5μg/ml fibronectin.

For live imaging experiments using a Nikon Ti-E microscope (Movies 1,3-5), cells were plated onto an 8-well imaging slide coated with 7.5μg/ml fibronectin, at the following densities: mCherry induction experiments: 3×10^4^ cells/well; mCherry downregulation experiments: 0-24h: 2×10^4^ cells/well; 24-48h: 10^4^ cells/well. For live imaging experiments using a PerkinElmer Opera Phenix Plus microscope (Movie 2, Fig. S16G), cells were plated onto a 96-well imaging plate coated with 7.5μg/ml fibronectin, at the following densities: mCherry induction experiments: 9.6×10^4^ cells/well; mCherry downregulation experiments: 0-24h: 2×10^4^ cells/well; 24-48h: 10^4^ cells/well. ESC culture medium was supplemented with 2μg/ml puromycin, 200μg/ml hygromycin B and 1X penicillin/streptomycin.

To test induction of 3xFlag-Neurog1 in STN receiver cells, CmGP1 sender cells and STN receiver cells were plated at a 9:1 ratio at a concentration of 2×10^5^ cells/well onto a 24mm flamed glass coverslip housed in a 6-well plate coated with 7.5μg/ml fibronectin, then fixed and stained after 48 hours.

### Synthetic patterning experiments

A 24mm glass coverslip housed in a 6-well plate was coated with 7.5μg/ml fibronectin, then allowed to air dry. When fully dry, forceps were used to place a culture insert 3-well silicon chamber on top of the coverslip; downward force was carefully exerted to secure it in place. 4×10^4^ cells were plated overnight in 70μl culture medium in each of the three wells. Sender cells were plated in the central well, and receiver cells were plated in the outside wells. 2ml culture medium were added outside of the 3-well insert in order to prevent evaporation. The next day, the 2ml culture medium outside of the 3-well insert were aspirated, and the 70μl in each of the three wells were carefully removed in order not to dislodge the 3-well insert. Each well was quickly washed with 70μl PBS to get rid of any remaining cells in suspension. Forceps were used to detach the 3-well insert from the glass coverslip, and 2.5ml culture medium were added to the well. Culture medium was replaced daily. Growth of cells into the gaps between wells were monitored daily; following contact between sender and receiver cells, cells were kept in culture for a further 24 hours (STC receivers + CmGP1 senders) or a further 96 hours (STN receivers + CmGP1 senders) prior to fixation and immunofluorescence. For live imaging of mCherry stripe experiments, the 3-well insert was placed in an Ibidi μ-Slide 4 Well instead of on a glass coverslip.

### Flow cytometry

Cells were washed in PBS, then detached from culture vessels with accutase. They were resuspended in ice-cold PBS+10% FCS, pelleted by spinning at 300g for 3 minutes, resuspended in ice-cold PBS+10% FCS+300nM DRAQ7 and placed on ice before analysing on a BD LSRFortessa flow cytometer. Forward and side-scatter width and amplitude were used to identify single cells in suspension, dead cells were excluded by gating on DRAQ7-negative cells, and tagBFP, GFP and mCherry/mKate2 expression were then analysed using V 450/50-A, B 530/30-A, Y/G 610/20-A laser/filters combinations respectively.

### Immunofluorescence

Cells were plated on flamed glass coverslips coated with 7.5μg/ml fibronectin and cultured as indicated in figure legends. Cells were washed with PBS, fixed in 4% formaldehyde in PBS for 20 minutes at room temperature, then washed three times in PBS for a total of 15 minutes. Cells were blocked overnight at 4°C in blocking solution (PBS+3% donkey serum+0.1% Triton X-100). Primary antibodies diluted in blocking solution were added for 3 hours at room temperature, the coverslips were washed 3 times in PBS for a total of 30 minutes, secondary antibodies diluted in blocking solution were added for 1 hour at room temperature, the coverslips were washed 3 times in PBS for a total of 30 minutes. The coverslips were then mounted onto glass slides in Prolong Gold mounting medium. For synthetic patterning experiments and chimaera stainings, antibodies were incubated overnight at 4°C or 37°C respectively to improve penetration. Blastocysts were imaged in PBS in an imaging chamber, and scoring of mCherry-HI cells was performed manually using chimaeras containing both sender and receiver cells as a reference. Post-implantation chimaeras were were dehydrated in methanol series in PBS/0.1% Triton X-100, clarified in 50% methanol/50% BABB (benzyl alcohol:benzyl benzoate 1:2 ratio), and transferred into 100% BABB before imaging. All imaging was performed on a Leica SP8 confocal microscope with a 40X immersion lens unless otherwise indicated.

### Live imaging

Cells to be imaged were allowed to adhere on culture vessels at room temperature for 15 minutes after plating, after which they were placed in a 37°C 5% CO_2_ humidified chamber and imaged. For Movies 1, 3-5, imaging was performed with a widefield Nikon Ti-E microscope, 20X lens, and Hamamatsu camera; images were taken at 10-minute intervals for 24 hours, and xy coordinates were saved. Following live imaging, cells were fixed and stained for fluorophore expression, and imaged at the previously saved xy coordinates (Movies 1,3). For Movie 2, Fig. S16G and Fig. S19, imaging was performed with a PerkinElmer Opera PhenixPlus microscope, 20X lens. Images were taken at 1-hour (Movie 2, Fig. S16G) or 90-minute (Fig. S19) intervals for 24 hours. Fully automated segmentation of tagBFP-positive nuclei, tracking, and quantification of fluorescent signal intensity in live imaging experiments was performed using the PerkinElmer Harmony software.

Filming of the morula to blastocyst transition was performed with a PerkinElmer Opera PhenixPlus confocal microscope. Sender and receiver cells were placed on opposite poles of morulae, and allowed to aggregate unperturbed for approximately 4 hours prior to imaging, in order to ensure strong binding of ES cells to morulae. They were then transferred to wells of an uncoated Ibidi μ-Slide Angiogenesis imaging slide and imaged. Aside from an initial frame, it was not possible to capture tagBFP signal within the time-lapse movies as embryos were vulnerable to repeated stimulation with ultraviolet light.

## ACKNOWLEDGEMENTS

We thank Leonardo Morsut and Wendell Lim for developing the original SynNotch system, Elise Cachat and Keisuke Kaji for advice and reagents, Luis Ángel Fernández for the pDisplay-EGFP-TM plasmid, Andras Nagy for the CAG-ϕC31 plasmid, Dónal O’Carroll and Pedro Moreira for assistance with generating chimaeric mice, Eve Moutaux for cloning initial versions of the RMCE constructs, Alexandre Veiga and Matthew French for performing preliminary experiments with the original SynNotch constructs. We thank Justyna Cholewa-Waclaw, Matthieu Vermeren and Charles Williams for assistance with time-lapse imaging, Fiona Rossi and Claire Cryer for flow cytometry support, and Theresa O’Connor, Helen Henderson and Marilyn Thomson for cell culture support. We are grateful to Leonardo Morsut and members of the Lowell and Wilson labs for helpful discussions.

