## Supplemental Information for "SyNPL: Synthetic Notch pluripotent cell lines to monitor and manipulate cell interactions *in vitro* and *in vivo*"

#### Supplementary information

consisting of

Supplementary Methods
Supplementary Figures S1-S21
Legends for Movies 1-6

### Supplementary Methods


**Design and characterisation of an "all-in-one" design for mCherry inducible receiver cells**

We generated a donor plasmid for recombination-mediated cassette exchange (RMCE) into EM35 cells, with a Pac (puromycin resistance) gene for positive selection of recombinant clones, and convenient restriction sites for the inclusion of several transcriptional units. We made use of this tool to generate ESC clonal lines containing the SynNotch receptor, the constitutive tagBFP transgene and the TRE-mCherry all within the *Rosa26* locus (Figs. S4-S6). We named these cells SNCB, for SynNotch-mCherry-tagBFP. The tagBFP and TRE-mCherry cassettes were either placed on the + strand (SNCB+, Fig. S4A) or on the - strand (SNCB-, Fig. S6A).

We screened 30 SNCB+ clones by analysing the levels of mCherry and tagBFP fluorescence in the absence of sender cells (Fig. S4B-E). Although all clones should be genetically identical, we observed variation between them: 2 clones expressed no mCherry nor tagBFP, suggesting they may have integrated only part of the construct, whereas the remaining 28 clones had broad variations in tagBFP expression, with a subset of tagBFP-high cells displaying low levels of mCherry expression (Fig. S4E). We selected two clones with low levels of mCherry leakiness (SNCB+4, SNCB+16) and co-cultured these with 4 different sender clones for 24 hours, a period of time shown to be sufficient for mCherry induction by SynNotch in L929 fibroblasts (Morsut et al., 2016) (Fig. S5A-L). The small subset of mCherry-leaky cells induced mCherry to higher levels, suggesting that the system was functional in this subset of cells, but the majority (>90%) of cells remained mCherry-negative (Figs. S5E,F,K,L). In order to exclude the unlikely possibility that lack of mCherry induction in receiver ESCs was due to lack of interaction with sender cells, we imaged SNCB+4 cells cultured alone or in the presence of CHmGMP19 sender cells for 24 hours. We did not observe mCherry expression in the absence of sender cells, but, as seen in the flow cytometry data, a proportion of cells did not express tagBFP either (Fig. S5M). We observed mCherry induction in a small number of cells following co-culture with sender cells, but most EGFP-negative cells in contact with EGFP-positive sender cells did not induce mCherry expression (Fig. S5M).

We then screened 7 SNCB- clones as above (Fig. S6B-E). The fluorescence distribution looked radically different from the SNCB+ clones, with only one clone (SNCB-6) displaying a clear tagBFP-positive subpopulation of cells (Fig. S6E). Co-culture with sender cells for 24h resulted in higher levels of mCherry expression in this subpopulation, which suggests the system is functional in these cells, but approximately half of the cells in this clone remained mCherry-negative (Fig. S6E).

Taken together, these results suggest that subsets of cells within SNCB+ and SNCB- clonal lines have the capability of functioning as SynNotch receiver cells. We therefore asked whether isolation and re-cloning of these cells may lead to the generation of functional SynNotch receiver lines.

We attempted to subclone cells from the SNCB+4 and SNCB-6 lines by sorting single cells based on their tagBFP and mCherry levels and generating new clonal lines. We selected cells which were tagBFP-high and mCherry-low in the absence of sender cells (+4BHCL, -6BHCL), tagBFP-high and mCherry-high in the presence of sender cells (+4BHCH, -6BHCH), and rare SNCB+4 cells which were tagBFP-medium and mCherry-positive in the presence of sender cells (+4BMC+) (Figs. S7A-C, S8A-C). After clonal expansion, a subset of cells in all clones lost tagBFP and mCherry expression, with overall tagBFP and mCherry expression patterns looking remarkably similar to those of the two parental lines. +4BHCH and -6BHCH cells had slightly better inducibility than the parental lines, but a significant proportion of cells lost tagBFP and mCherry expression (Figs. S7D-F, S8D,E). This suggests that sorting and subcloning of SNCB+4 and SNCB-6 cells did not result in the generation of functional receiver ESCs.

We next asked whether lack of mCherry induction in receiver cells could be ascribable to the SynNotch receptor construct not being expressed by all cells. We stained wild-type, SNCB+4 and CHmGMP19 cells for Myc expression (Fig. S9); a Myc tag is present on both the SynNotch receptor construct and the extracellular EGFP in CHmGMP19 cells (Figs. 2B,J, S4A). We found that although all SNCB+4 cells expressed Myc above background level, its pattern of expression only appeared to be consistent with cell membrane localisation in a subset of cells. Furthermore, the levels of expression were significantly lower than those of Myc-EGFP in CHmGMP19 sender cells (Fig. S9). .

In summary, receiver cell lines harbouring all three transcriptional units required for SynNotch receiver function within the Rosa26 landing pad do not exhibit uniform behaviour, displaying variable levels of tagBFP, variable inducibility of mCherry, and low levels of the SynNotch receptor. Despite this, mCherry is induced by subpopulations of tagBFP-positive receiver cells in response to interaction with EGFP-positive sender cells. These results suggest that the SynNotch receptor construct and TRE-mCherry cassette can function as expected in ESCs, but that further optimisation is required to obtain a reliable contact-reporting system. We have included this suboptimal strategy in this report in order to inform researchers who wish to establish SynNotch technology in other experimental models of interest.

### Supplementary Methods

**Generation and characterisation of synthetic stripe pattern of mCherry expression**

In order to generate a stripe pattern of mCherry expression, $4x10^4$ sender and STC receiver cells were plated in adjacent wells of a 3-well cell culture insert and left to attach and spread overnight. The next morning, wells were washed with PBS to remove any floating cells, and the insert was carefully removed to avoid detaching clumps of cells. The smaller the culture vessel the insert was housed in, the harder this became, with clumps of cells visibly detaching in small multiwell plates. Examples are displayed in Fig. S19A. Additional wash steps were used in these circumstances to reduce the risk of clumps settling down onto different cells and inducing mCherry expression before a linear border was formed (e.g. sender cells landing in the middle of the receiver "domain" or vice versa). This phenomenon could not always be prevented, and where possible we would recommend housing the insert onto a 24cm glass coverslip in a 6-well plate to reduce the risk of this process occuring.

Following insert removal, the expansion of sender and receiver cells was monitored in order to identify the time of initial contact. This varied between experiments, potentially as a result of leftover glue from the insert in the gap between sender and receiver cells affecting cell proliferation and migration. Initial contact was made approximately 48 hours after insert removal (Fig. S19A). Formation of a stripe along the entire length of the border was evident by 24 hours after initial contact (Fig. S19A), in line with the kinetics of mCherry upregulation in non-patterned culture (Fig. 4).

Timelapse imaging of stripe formation is complicated by the thickness and size of the structure to be imaged, which in our setup led to sample bleaching (Fig. S19B), making it inappropriate to draw fluorescence intensity-based conclusions.

Over time, the mCherry signal changes from a narrow signal at the point of contact to a diffuse gradient into the receiver cell domain (Fig. S19A,B). This is caused by several factors:

1. Sender cells can infiltrate into the receiver cell domain at the bottom of the dish, activating mCherry expression in patches of cells beyond the linear border (Fig. S19B).
2. Sender cells can migrate on top of receiver cells, activating mCherry induction beyond the linear border (Fig. S19C).
3. Activated receiver cells can divide perpendicularly to the linear border, contributing to diffusion of mCherry signal (Fig. S19B, yellow arrowheads).

#### FIGURE S1

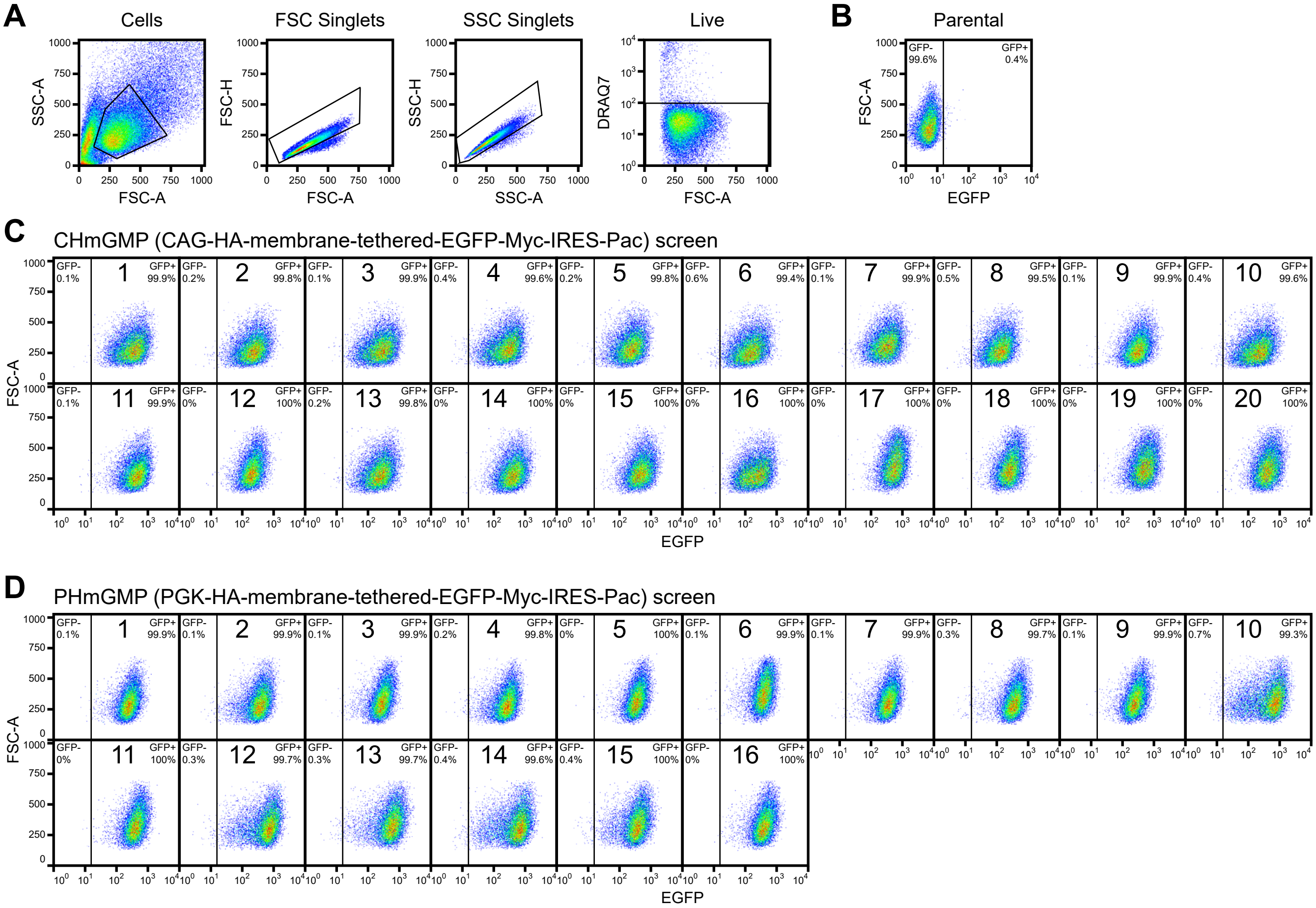


**Figure S1. Screening of HA- and Myc-tagged EGFP clonal sender ESC lines.**
(A) Flow cytometry gating strategy to analyse single live cells, making use of forward scatter (FSC) and side scatter (SSC) height (H) and amplitude (A), and of the cell-impermeable DRAQ7 nuclear counterstain. (B) EGFP fluorescence distribution in parental wild-type cells. (C-D) EGFP fluorescence distribution in (C) CHmGMP and (D) PHmGMP clones. Percentages of EGFP-positive and -negative cells are indicated in figure. 15000 cells were analysed for each sample in (B-D). All units of measurement are arbitrary fluorescence units (A.F.U.). n=1.

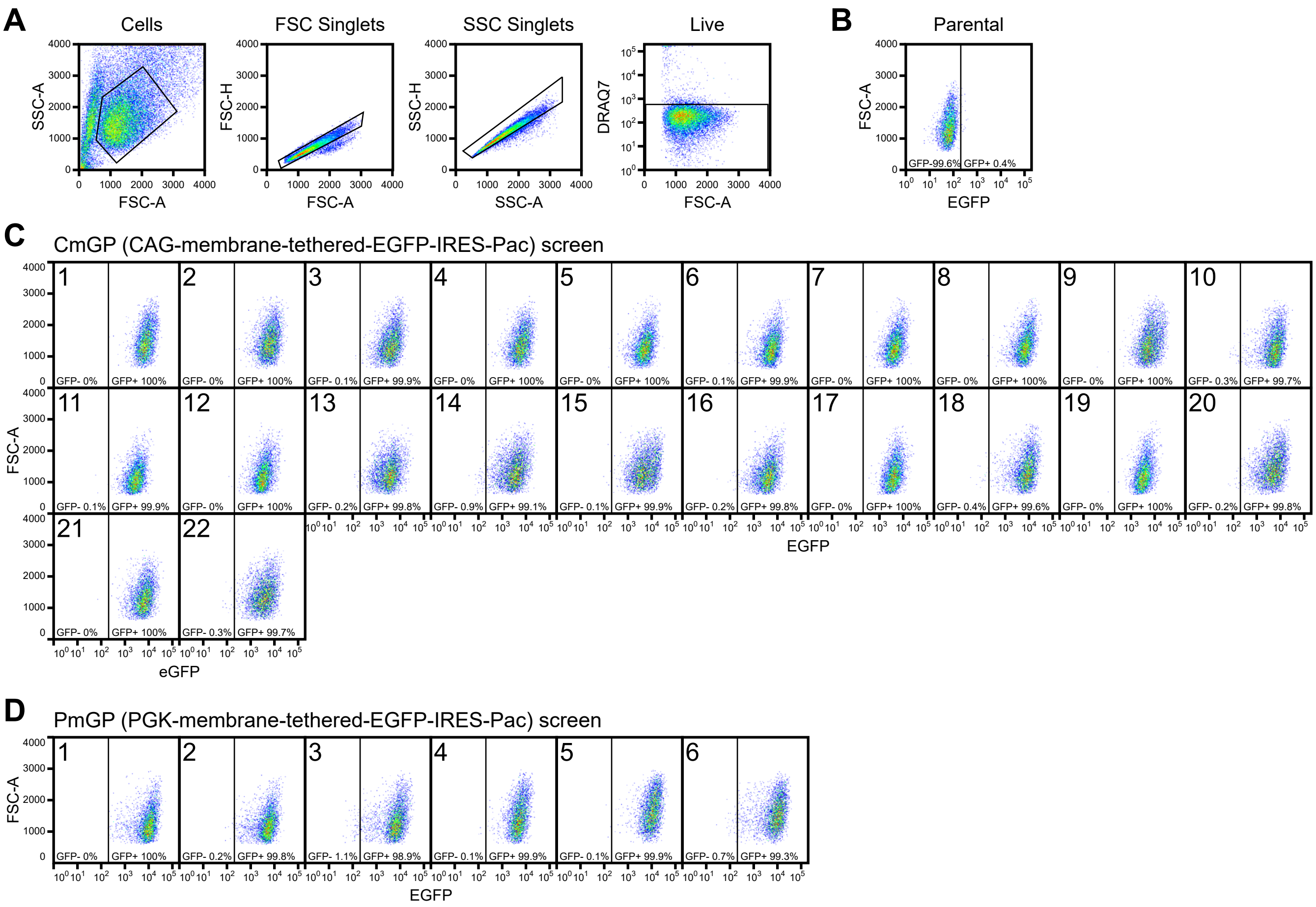


**Figure S2. Screening of untagged EGFP clonal sender ESC lines.**
(A) Flow cytometry gating strategy to analyse single live cells. (B) EGFP fluorescence distribution in parental wild-type cells. (C-D) EGFP fluorescence distribution in (C) CmGP and (D) PmGP clones. Percentages of EGFP-positive and -negative cells are indicated in figure. 5000 cells were analysed for each sample in (B-D). All units of measurement are arbitrary fluorescence units (A.F.U.), and values are not directly comparable to those in Fig. S1. n=1.

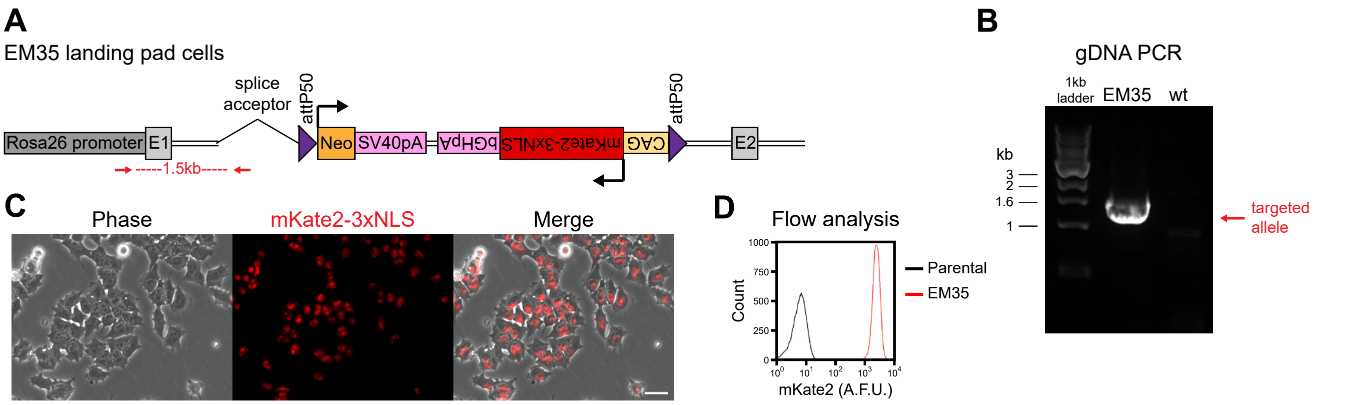


**Figure S3. Generation of a safe harbour site landing pad master ESC line.**
(A) Diagram of targeted Rosa26 allele in EM35 landing pad RMCE clonal ESC line. Locations of primers for testing of correct targeting event are shown as red arrows. Abbreviations: E1: exon 1; E2: exon 2. (B) Genomic DNA PCR validation of correct landing pad insertion at the *Rosa26* locus, using the primers shown in (A). Expected band sizes: correct targeting event: 1534bp; wild-type: no amplification. Abbreviations: gDNA: genomic DNA; wt: parental wild-type. (C) Phase-contrast and mKate2-3xNLS images of live EM35 ESCs, showing nuclear mKate2 signal in all cells. Scale bar: 30μm. (D) Flow cytometry analysis of mKate2-3xNLS expression in parental wild-type and EM35 cells. 20000 cells are displayed for each sample. Data from a single experiment, representative of five biological replicates.

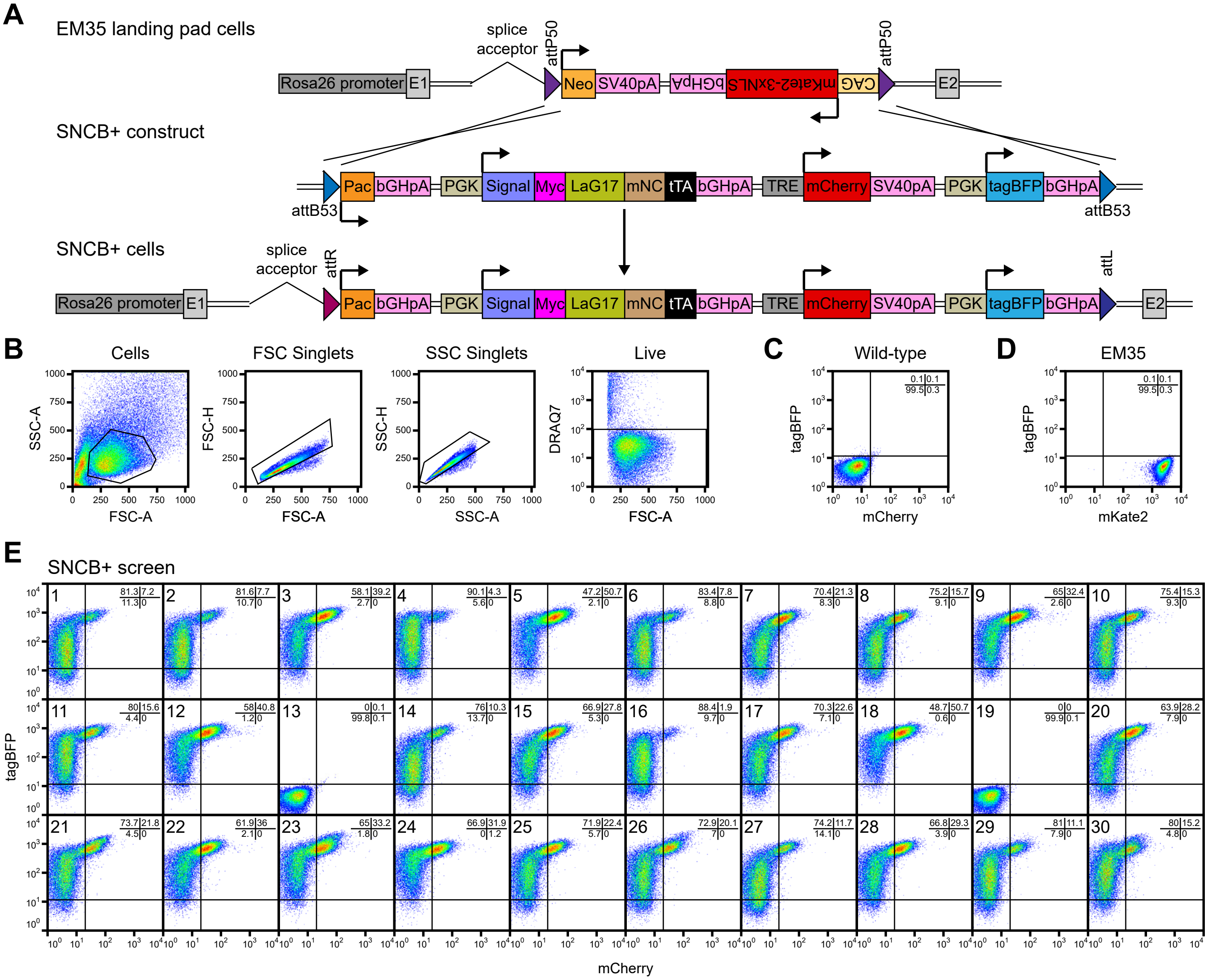


**Figure S4. Generation and screening of SNCB+ clonal receiver ESC lines.**
(A) Strategy to replace Neo-mKate2 cassette with SNCB+ cassette in EM35 landing pad ESCs through φC31 integrase-mediated RMCE. (B) Flow cytometry gating strategy to analyse single live cells. (C) tagBFP and mCherry fluorescence distribution in wild-type ESCs. (D) tagBFP and mKate2 fluorescence distribution in parental EM35 landing pad ESCs. The same laser/filter combinations were used to detect both mKate2 and mCherry fluorescence. (E) tagBFP and mCherry fluorescence distribution in SNCB+ clonal ESC lines cultured in the absence of sender cells. Percentages of cells in each quadrant are indicated in figure. 35000 cells were analysed for each sample in (C-E). All units of measurement are arbitrary fluorescence units (A.F.U.). n=1.

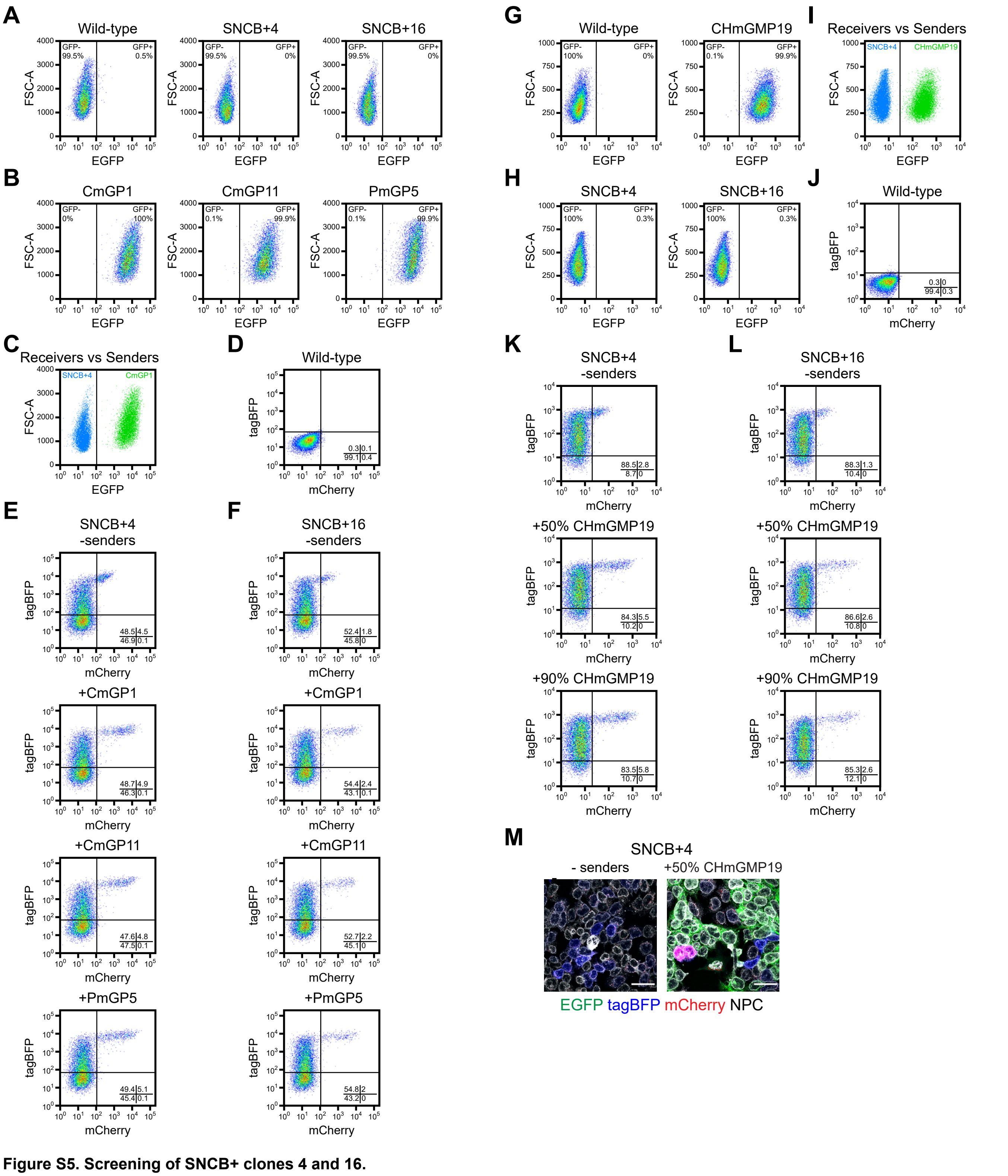


**Figure S5. Screening of SNCB+ clones 4 and 16.**
(A-B) EGFP fluorescence distribution in (A) wild-type, SNCB+4, SNCB+16 receiver cells and (B) CmGP1, CmGP11, PmGP5 sender cells. (C) Comparison of EGFP fluorescence distribution in SNCB+4 receiver cells and CmGP1 sender cells. The two cell populations can be separated in co-culture experiments on the basis of EGFP expression. (D) tagBFP and mCherry fluorescence distributions in wild-type cells. (E-F) tagBFP and mCherry fluorescence distributions in (E) SNCB+4 and (F) SNCB+16 receiver ESCs cultured alone or in the presence of CmGP1, CmGP11, PmGP5 sender ESCs for 24 hours. All units of measurement are arbitrary fluorescence units (A.F.U.). (G-H) EGFP fluorescence distribution in (G) wild-type, CHmGMP19 sender cells and (H) SNCB+ clones 4 and 16 receiver cells. (I) Comparison of EGFP fluorescence distribution in SNCB+4 receiver cells and CHmGMP19 sender cells. The two cell populations can be separated in co-culture experiments on the basis of EGFP expression. (J) tagBFP and mCherry fluorescence distributions in wild-type cells. (K-L) tagBFP and mCherry fluorescence distributions in (K) SNCB+4 and (L) SNCB+16 receiver ESCs cultured alone or in the presence of CHmGMP19 sender ESCs at 1:1 and 9:1 sender:receiver cell ratios for 24 hours. Experiments in panels (A-F) were carried out separately from those in panels (G-L) and fluorescence intensity values are not directly comparable. 15000 cells are displayed in plots in (D-F, G,H, J-L), 15000 cells/sample are displayed in plot in (I), 10000 cells are displayed in plots in (A,B), 10000 cells/sample are displayed in plot in (C). (M) Immunofluorescence of SNCB+4 cells cultured alone or in the presence of CHmGMP19 sender cells for 24 hours (1:1 sender:receiver cell ratio). Scale bar: 30μm. n=1.

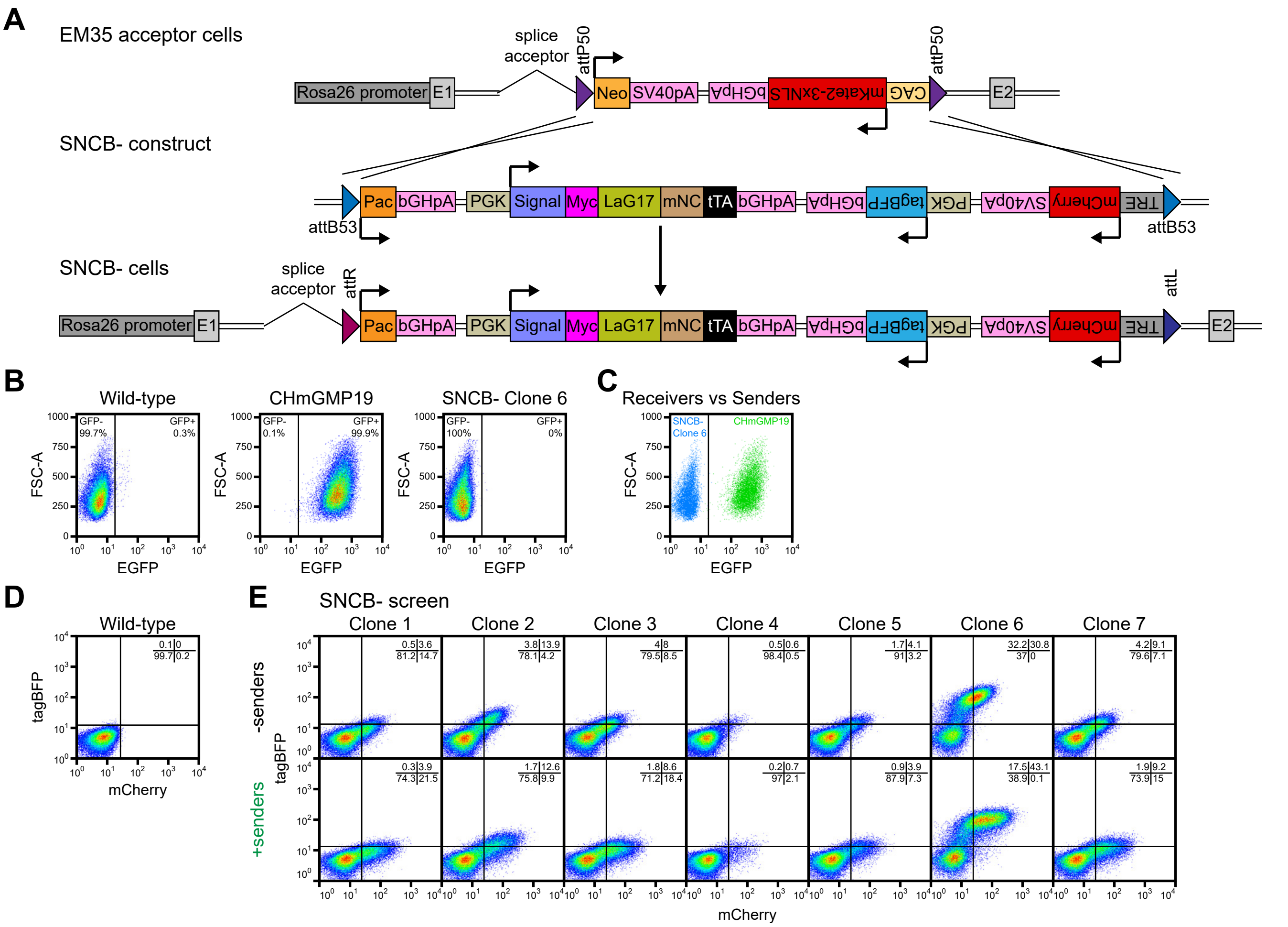


**Figure S6. Generation and screening of SNCB- clonal receiver ESC lines.**
(A) Strategy to replace Neo-mKate2 cassette with SNCB- cassette in EM35 landing pad ESCs through φC31 integrase-mediated RMCE. (B) EGFP fluorescence distribution in wild-type, CHmGMP19 sender cells and SNCB- clone 6 receiver cells. (C) Comparison of EGFP fluorescence distribution in SNCB- clone 6 receiver cells and CHmGMP19 sender cells. The two cell populations can be separated in co-culture experiments on the basis of EGFP expression. (D) tagBFP and mCherry fluorescence distribution in wild-type ESCs. (E) tagBFP and mCherry fluorescence distribution in SNCB- clonal ESC lines cultured alone or in the presence of CHmGMP19 sender cells for 24 hours. Percentages of cells in each quadrant are indicated in figure. 35000 cells are displayed in plots in (B), 35000 cells/sample are displayed in plot in (C), 50000 cells are displayed in plots in (D-E). All units of measurement are arbitrary fluorescence units (A.F.U.). n=1.

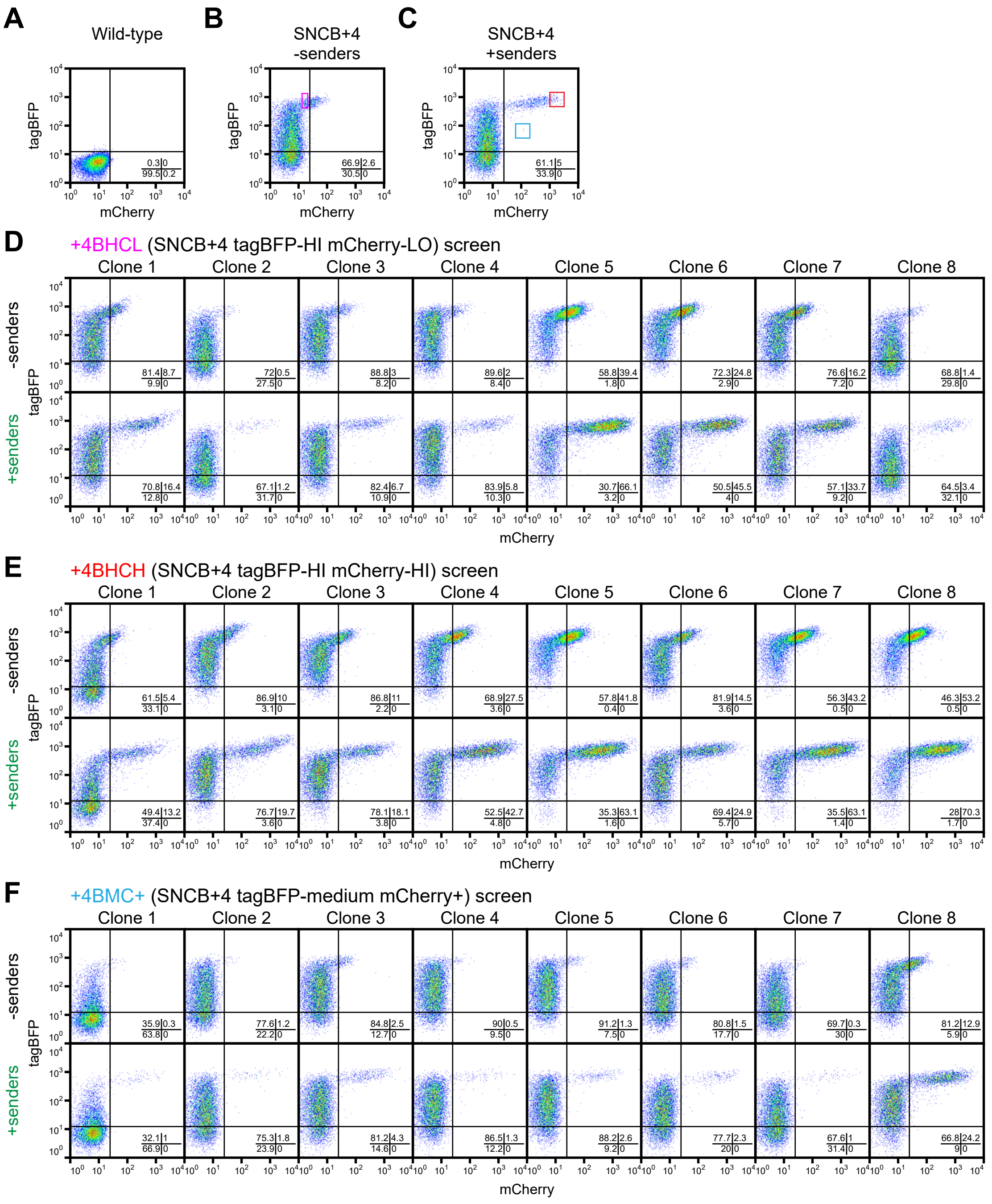


**Figure S7. Derivation of clonal ESC lines through sorting of SNCB+4 ESCs.**
(A) tagBFP and mCherry fluorescence distribution in wild-type ESCs. (B) tagBFP and mCherry fluorescence distribution in SNCB+4 ESCs cultured in the absence of sender cells. Sorting gate for derivation of SNCB+4 tagBFP-HI mCherry-LO (+4BHCL) clonal sorted lines is shown in magenta. (C) tagBFP and mCherry fluorescence distribution in SNCB+4 ESCs co-cultured with CHmGMP19 sender cells for 24 hours. Sorting gate for derivation of SNCB+4 tagBFP-HI mCherry-HI (+4BHCH) clonal sorted lines is shown in red, sorting gate for derivation of SNCB+4 tagBFP-Medium mCherry+ (+4BMC+) clonal sorted lines is shown in blue. (D-F) tagBFP and mCherry fluorescence distribution in (D) +4BHCL, (E) +4BHCH and (F) +4BMC+ receiver ESCs cultured alone or in the presence of CHmGMP19 sender cells for 24 hours. Percentages of cells in each quadrant are indicated in figure. 15000 cells are displayed in plots in (A-C), 10000 cells are displayed in plots in (D-F). All units of measurement are arbitrary fluorescence units (A.F.U.). n=1.

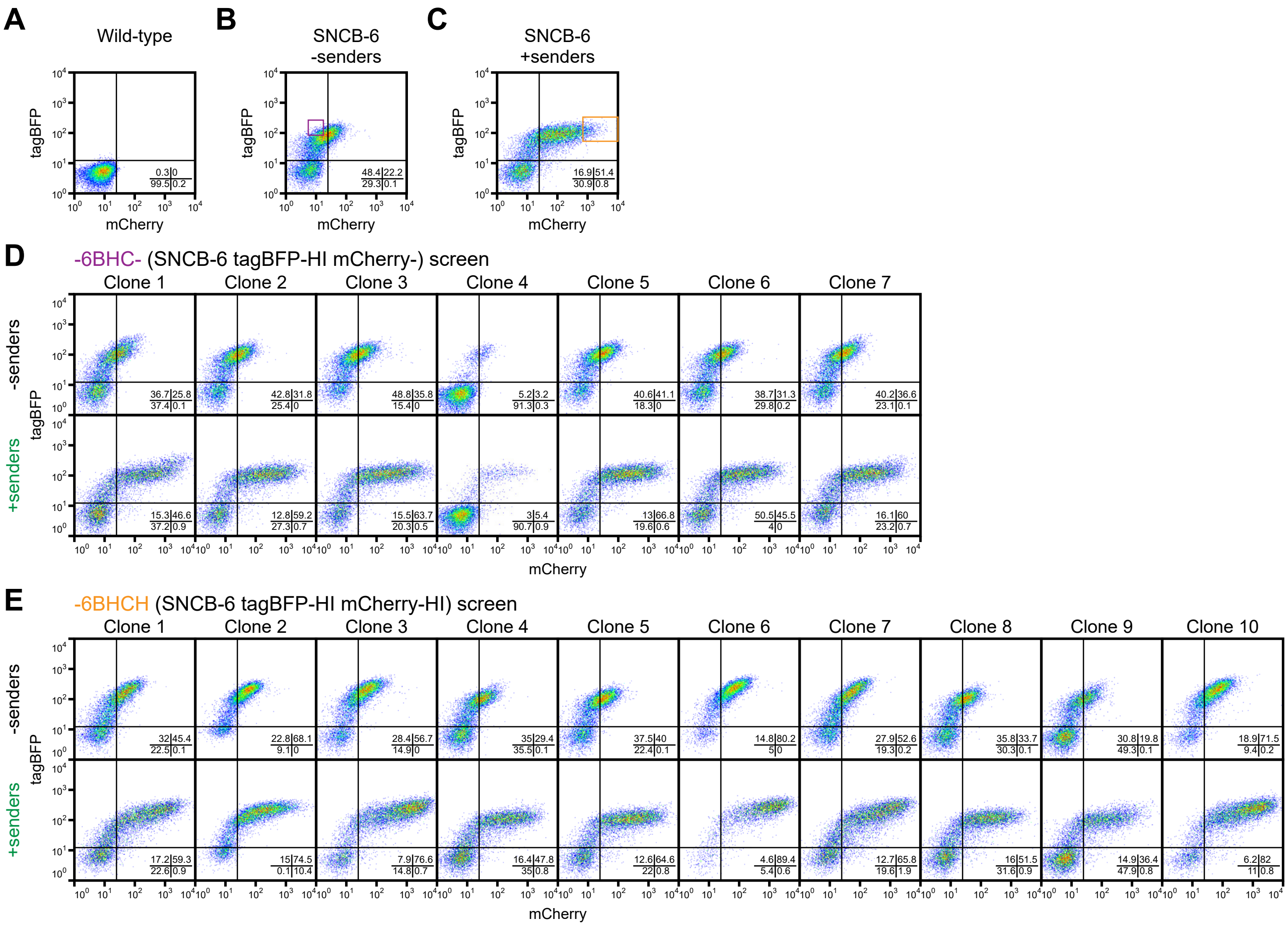

**Figure S8. Derivation of clonal ESC lines through sorting of SNCB-6 ESCs.**
(A) tagBFP and mCherry fluorescence distribution in wild-type ESCs. (B) tagBFP and mCherry fluorescence distribution in SNCB-6 ESCs cultured in the absence of sender cells. Sorting gate for derivation of SNCB-6 tagBFP-HI mCherry- (-6BHC-) clonal sorted lines is shown in purple. (C) tagBFP and mCherry fluorescence distribution in SNCB-6 ESCs co-cultured with CHmGMP19 sender cells for 24 hours. Sorting gate for derivation of SNCB-6 tagBFP-HI mCherry-HI (-6BHCH) clonal sorted lines is shown in orange. (D-E) tagBFP and mCherry fluorescence distribution in (D) -6BHC- and (E) -6BHCH receiver ESCs cultured alone or in the presence of CHmGMP19 sender cells for 24 hours. Percentages of cells in each quadrant are indicated in figure. 15000 cells are displayed in plots in (A-C), 10000 cells are displayed in plots in (D-E). All units of measurement are arbitrary fluorescence units (A.F.U.). n=1.

#### FIGURE S9

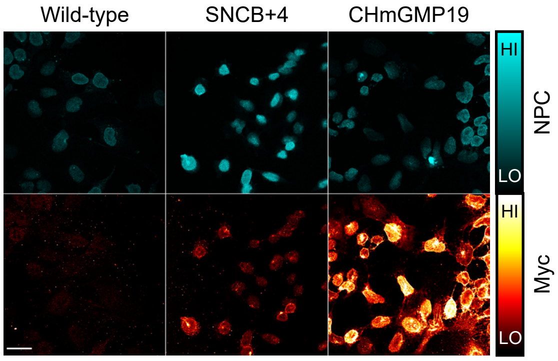


**Figure S9. The SynNotch receptor is expressed at low levels in SNCB+4 ESCs.**
Immunofluorescence of wild-type, SNCB+4 and CHmGMP19 cells. Colour lookup tables for NPC and Myc are displayed on the right of the images. Scale bar: 30μm.

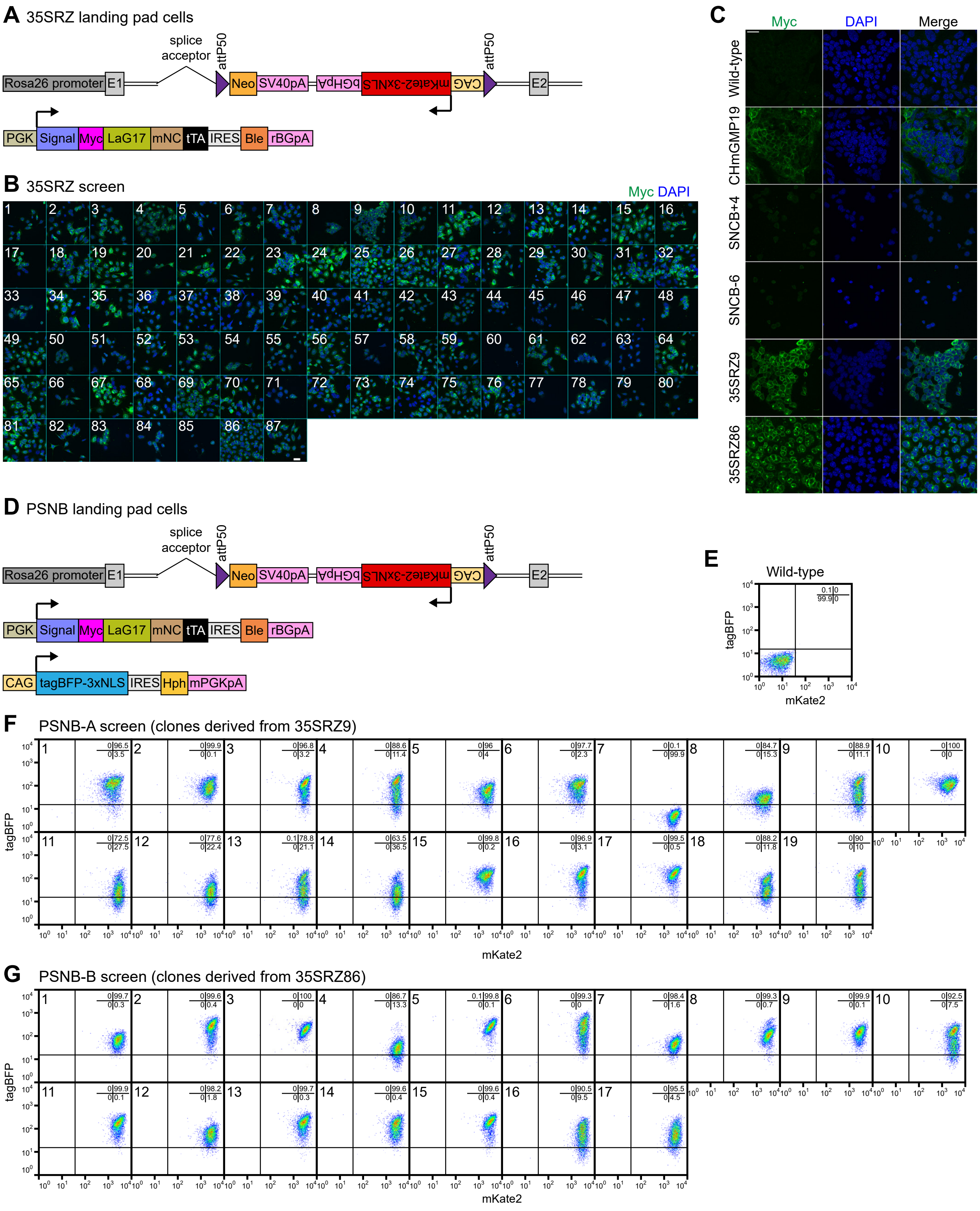


**Figure S10. Generation of 35SRZ and PSNB clonal landing pad ESC lines.**
(A) Summary of transgenes stably integrated into the genome of 35SRZ clonal ESC lines. (B) Myc immunofluorescence in 87 35SRZ clonal ESC lines. Nuclei are counterstained with DAPI. Scale bar: 30µm. (C) Myc immunofluorescence in wild-type, CHmGMP19 sender cells, SNCB+4, SNCB-6 receiver cells, 35SRZ clones 9 and 86 landing pad cells. The images for 35SRZ clones 9 and 86 are magnified versions of those in panel (B). (D) Summary of transgenes stably integrated into the genome of PSNB clonal ESC lines. (E) tagBFP and mKate2 fluorescence distribution in wild-type ESCs. (F) tagBFP and mKate2 fluorescence distribution in 19 PSNB-A clonal ESC lines. These lines were derived from 35SRZ clone 9 ESCs. (G) tagBFP and mKate2 fluorescence distribution in 17 PNSB-B clonal ESC lines. These lines were derived from 35SRZ clone 86 ESCs. Percentages of cells in each quadrant are indicated in figure. 6500 cells are displayed in flow cytometry plots. All units of measurement are arbitrary fluorescence units (A.F.U.). n=1.

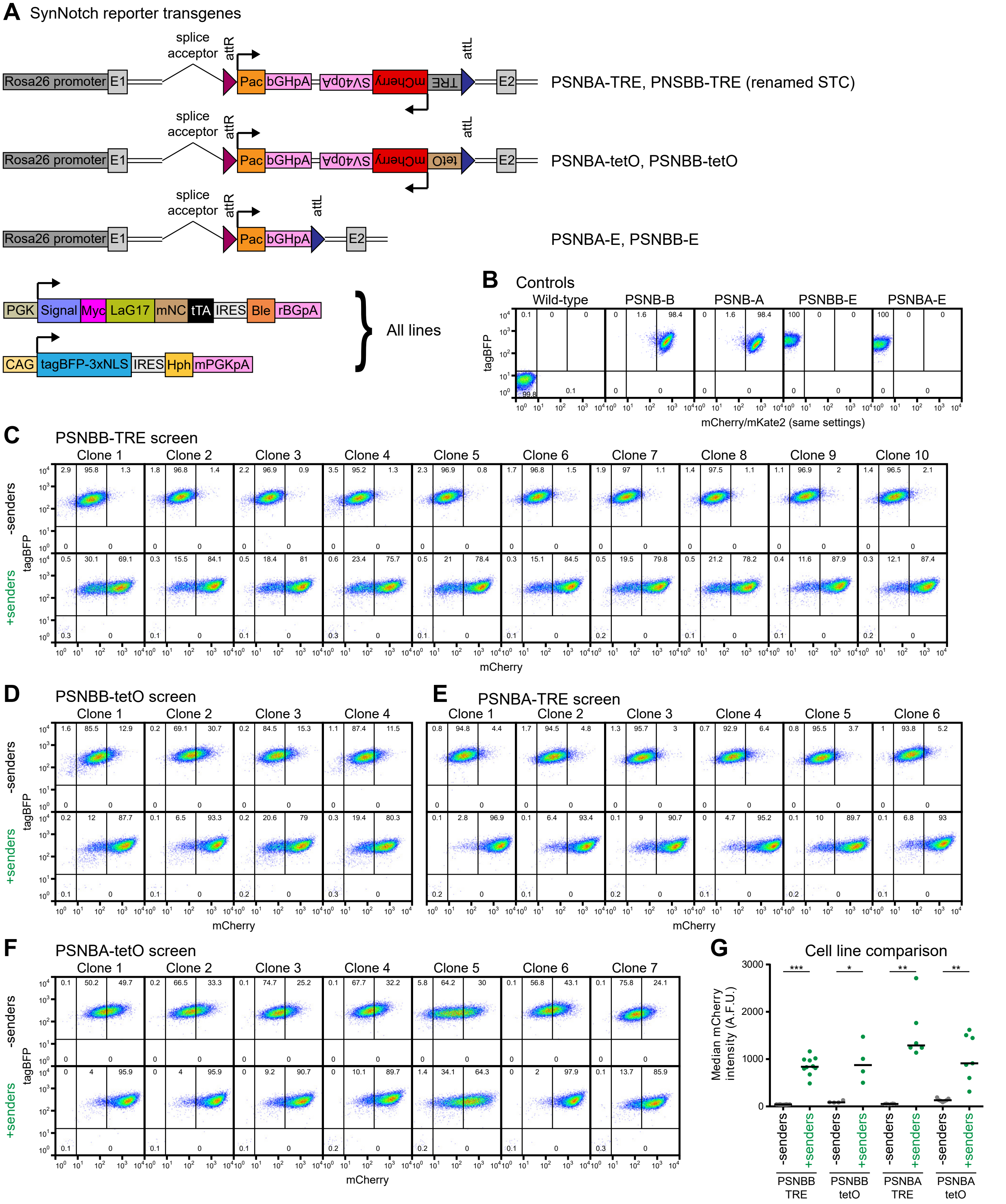


**Figure S11. Generation and screening of PSNB-TRE (STC), PSNB-tetO and PSNB-E clonal receiver ESC lines.**
(A) Summary of transgenes stably integrated into the genome of PSNB-TRE (STC), PSNB-tetO and PSNB-E clonal ESC lines. (B) tagBFP and mCherry/mKate2 fluorescence distribution in wild-type, PSNB and PSNB-E ESCs. (C-F) tagBFP and mCherry fluorescence distribution in (C) PSNBB-TRE, (D) PSNBB-tetO, (E) PSNBA-TRE and (F) PSNBA-tetO clonal ESC lines cultured alone or in the presence of CHmGMP19 sender cells for 24 hours. Percentages of cells in each quadrant are indicated in figure. 15000 cells are displayed in flow cytometry plots. All units of measurement are arbitrary fluorescence units (A.F.U.). n=1. (G) Comparison of median mCherry levels in PNSBB-TRE, PNSBB-tetO, PSNBA-TRE and PSNB-tetO cell lines. Each dot represents a different clone. Horizontal bar: median. Paired t-test p-values: *<0.05, **<0.01, ***<0.001.

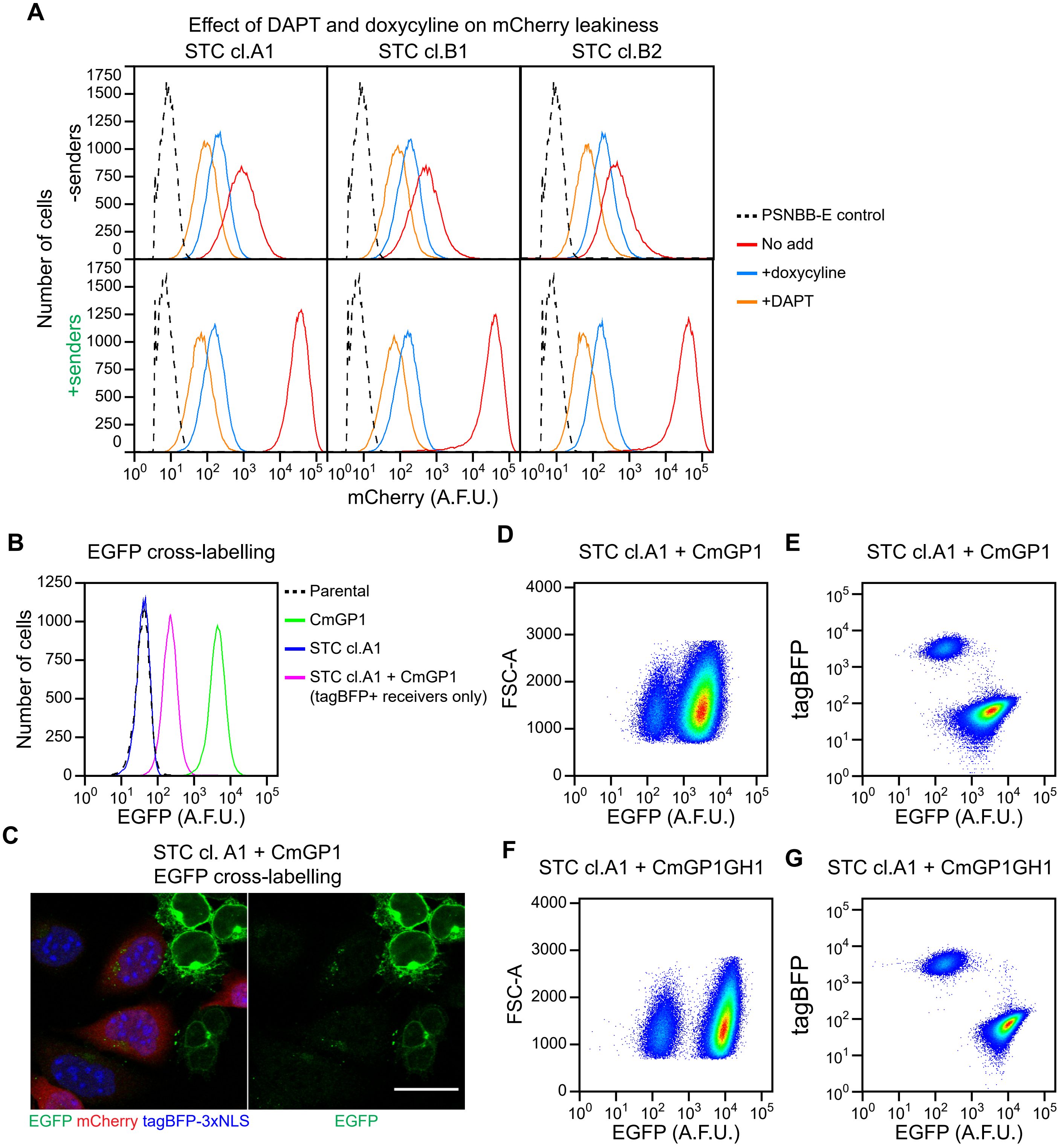


**Figure S12. Characterisation of STC receiver cells.**
(A) Flow cytometry analysis of mCherry expression in tagBFP+ EGFP- STC receiver clones A1, B1 and B2 cultured alone or in the presence of CmGP1 sender cells at a 9:1 sender:receiver ratio for 48 hours. Cells were cultured in the presence of 100µM DAPT, 1µg/ml doxycyline, or in the absence of both as indicated. PSNBB-E cells are included as a negative control. 40000 cells were analysed for each sample. (B) Flow cytometry analysis of EGFP expression in wild-type cells, CmGP1 sender cells, STC cl.A1 receiver cells cultured alone, and tagBFP+ STC cl.A1 receiver cells co-cultured with CmGP1 sender cells for 48 hours. 25000 cells were analysed for each sample. (C) Immunofluorescence analysis showing punctuate EGFP staining within STC cl.A1 receiver cells co-cultured with CmGP1 sender cells for 24h hours. Scale bar: 30µm. (D-G) Flow cytometry analysis of STC clone A1 receiver cells co-cultured with (D,E) CmGP1 or (F,G) CmGP1GH1 sender cells at a 9:1 sender:receiver ratio for 48 hours. 20000 cells were analysed for each sample. A.F.U. = arbitrary fluorescence units. All flow data comes from single experiments, representative of three biological replicates.

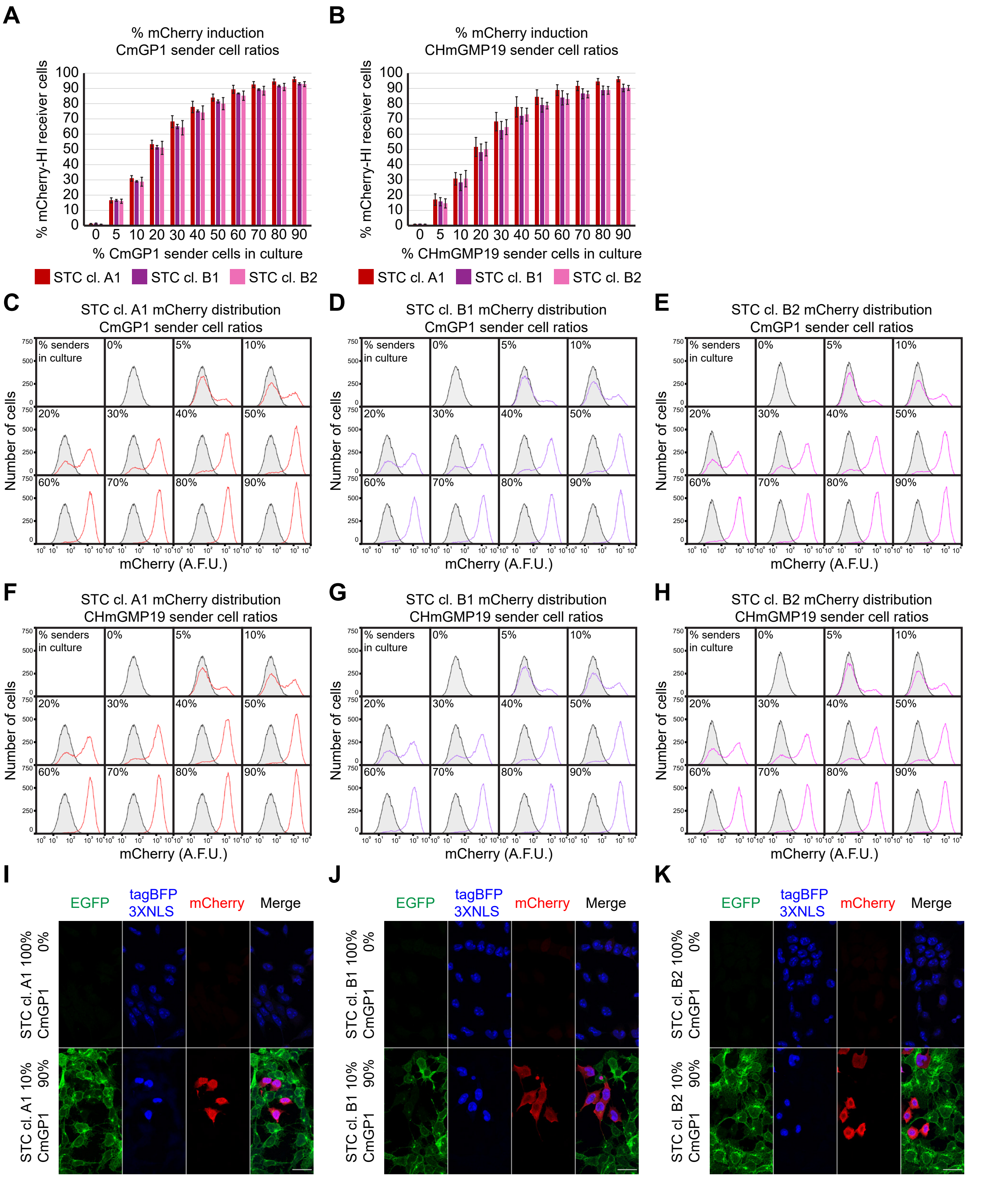


**Figure S13. Effect of varying sender:receiver cell ratios on mCherry induction in STC receiver cells.**
(A-B) Percentage of mCherry-HI STC receiver cells following 24 hours of co-culture with (A) CmGP1 or (B) CHmGMP19 sender cells at varying sender:receiver cell ratios. Data presented as mean ± standard deviation of three independent experiments. A minimum of 16000 cells were analysed for each sample. (C-H) Distribution of mCherry fluorescence in (C,F) STC clone A1, (D,G) STC clone B1 and (E,H) STC clone B2 receiver cells following 24 hours of co-culture with (C-E) CmGP1 or (F-H) CHmGMP19 sender cells at varying sender:receiver cell ratios. Data from a single experiment, representative of three biological replicates. STC clone A1, B1 and B2 cells cultured alone ("0%") are displayed as a shaded black histogram in all panels in (C-H). 23000 cells were analysed for each sample. A.F.U.: arbitrary fluorescence units. (I-K) Immunofluorescence of (I) STC clone A1, (J) STC clone B1 and (K) STC clone B2 receiver cells cultured alone or in the presence of CmGP1 sender cells (9:1 sender:receiver cell ratio). Scale bar: 30μm.

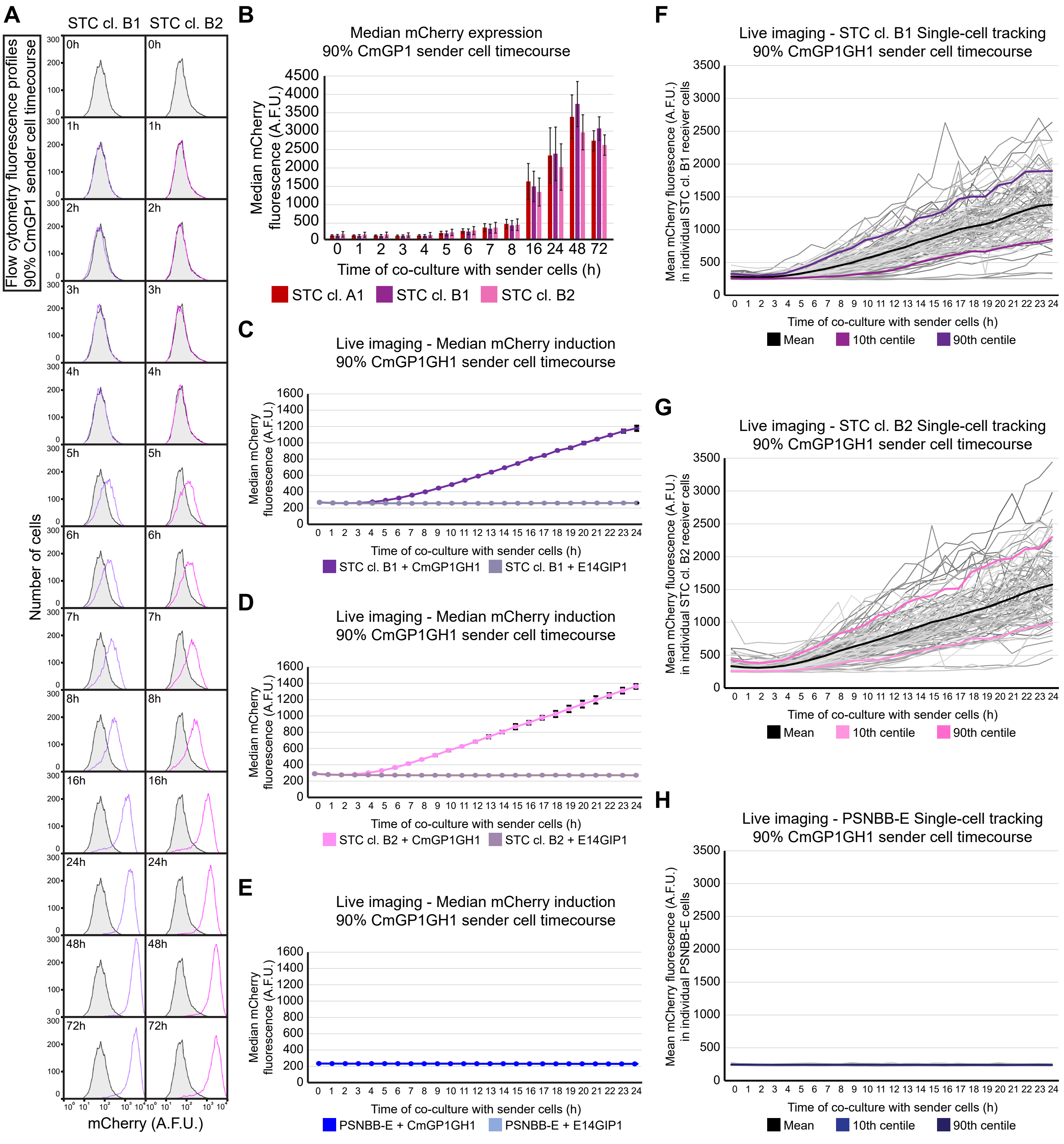


**Figure S14. Kinetics of mCherry induction in STC clones B1 and B2.**
(B) Flow cytometry analysis of distribution of mCherry fluorescence in STC clones B1 and B2 following co-culture with CmGP1 sender cells for the indicated amount of time (9:1 sender:receiver cell ratio). Data from a single experiment, representative of three biological replicates. Receiver cells cultured alone ("0h") are displayed as a shaded black histogram in all panels. 10000 cells were analysed for each sample. (B) Flow cytometry analysis of median mCherry expression in STC receiver cells following co-culture with CmGP1 sender cells for the indicated amount of time (9:1 sender:receiver cell ratio). Data presented as mean ± standard deviation of three independent experiments. A minimum of 8000 cells were analysed for each sample. (C-E) Quantification of live imaging: median mCherry fluorescence intensity in (C) STC clone B1 receiver cells, (D) STC clone B2 receiver cells, (E) PSNBB-E control cells following co-culture with CmGP1GH1 or E14GIP1 cells for the indicated amount of time (9:1 sender:receiver cell ratio). Average of 3 biological replicates, 10 fields of view/replicate, minimum of 446 cells/replicate/timepoint. Error bars: standard deviation. (F-H) Mean mCherry fluorescence intensity in individual (F) STC clone B1, (G) STC clone B2, (H) PSNBB-E cells tracked for 24 hours whilst in co-culture with CmGP1GH1 sender cells (9:1 sender:receiver ratio). Tracks are displayed for 33 cells for each of 3 biological replicates (99 cells total). Mean, 10th, and 90th centile tracks are also displayed. A.F.U.: arbitrary fluorescence units. All data in this figure were acquired at the same time as those in Figure 4, and are therefore directly comparable.

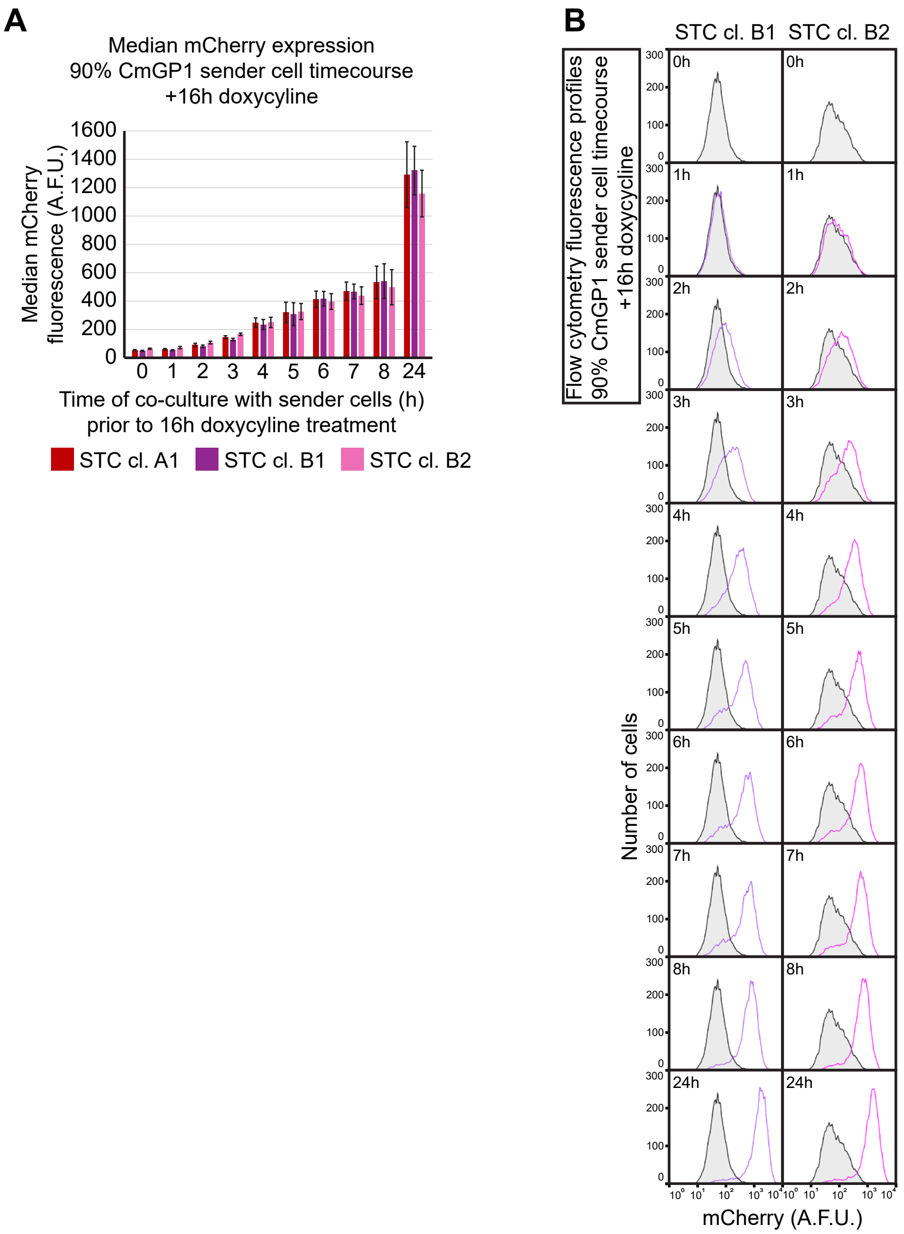


**Figure S15. Characterisation of minimal contact time required for mCherry induction in STC clones B1 and B2.**
(A) Median mCherry expression in STC receiver cells following co-culture with CmGP1 sender cells for the indicated amount of time and 16 hours doxycycline treatment (9:1 sender:receiver cell ratio). Data presented as mean ± standard deviation of three independent experiments. A minimum of 8000 cells were analysed for each sample. (B) Distribution of mCherry fluorescence in STC clones B1 and B2 following co-culture with CmGP1 sender cells for the indicated amount of time and 16 hours doxycycline treatment (9:1 sender:receiver cell ratio). Data from a single experiment, representative of three biological replicates. STC clone B1 and B2 cells plated with CmGP1 sender cells in doxycycline-containing medium for 16 hours (“0h”) are displayed as shaded black histograms in all panels. 10000 cells were analysed for each sample. A.F.U.: arbitrary fluorescence units. All data in this figure were acquired at the same time as those in Figure 5, and are therefore directly comparable.

**FIGURE S16**



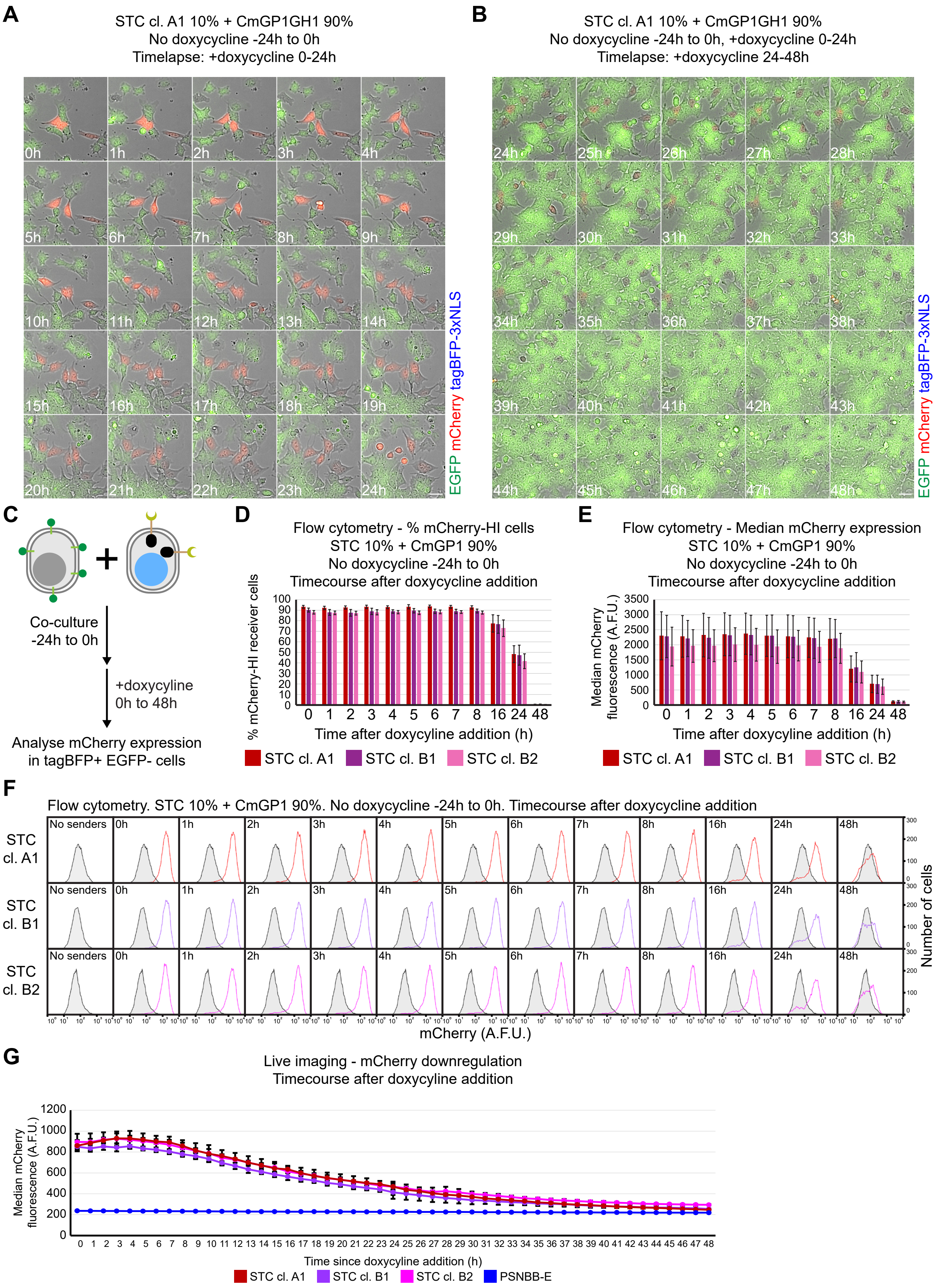
A
STC cl. A1 10% + CmGP1GH1 90%
No doxycycline -24h to 0h
Timelapse: +doxycycline 0-24h
0h
1h
2h
3h
4h
5h
6h
7h
8h
9h
10h
11h
12h
13h
14h
15h
16h
17h
18h
19h
20h
21h
22h
23h
24h
EGFP mCherry tagBFP-3xNLS
B
STC cl. A1 10% + CmGP1GH1 90%
No doxycycline -24h to 0h, +doxycycline 0-24h
Timelapse: +doxycycline 24-48h
24h
25h
26h
27h
28h
29h
30h
31h
32h
33h
34h
35h
36h
37h
38h
39h
40h
41h
42h
43h
44h
45h
46h
47h
48h
EGFP mCherry tagBFP-3xNLS
C
Co-culture
-24h to 0h
+doxycyline
0h to 48h
Analyse mCherry expression
in tagBFP+ EGFP- cells
D
Flow cytometry - % mCherry-HI cells
STC 10% + CmGP1 90%
No doxycycline -24h to 0h
Timecourse after doxycycline addition
% mCherry-HI receiver cells
Time after doxycycline addition (h)
STC cl. A1
STC cl. B1
STC cl. B2
E
Flow cytometry - Median mCherry expression
STC 10% + CmGP1 90%
No doxycycline -24h to 0h
Timecourse after doxycycline addition
Median mCherry fluorescence (A.F.U.)
Time after doxycyline addition (h)
STC cl. A1
STC cl. B1
STC cl. B2
F
Flow cytometry. STC 10% + CmGP1 90%. No doxycycline -24h to 0h. Timecourse after doxycycline addition
STC cl. A1
STC cl. B1
STC cl. B2
No senders
0h
1h
2h
3h
4h
5h
6h
7h
8h
16h
24h
48h
Number of cells
mCherry (A.F.U.)
G
Live imaging - mCherry downregulation
Timecourse after doxycycline addition
Median mCherry fluorescence (A.F.U.)
Time since doxycycline addition (h)
STC cl. A1
STC cl. B1
STC cl. B2
PSNBB-E

**Figure S16. Kinetics of mCherry downregulation in STC receiver cells.**
(A-B) Stills from Movies 4 and 5 displaying mCherry and EGFP expression in STC clone A1 receiver cells co-cultured with CmGP1GH1 sender cells for 24 hours (9:1 sender:receiver cell ratio), prior to treatment with 1μg/ml doxycycline. Filming was carried out between (A) 0 and 24 hours after doxycycline addition or (B) 24 and 48 hours after doxycycline addition. Scale bar: 30μm. (C) Experimental setup to analyse kinetics of mCherry downregulation in STC receiver cells by flow cytometry. Following sender:receiver cell co-culture for 24 hours, 1μg/ml doxycycline was added to the culture medium for 0-48 hours in order to inhibit tTA-mediated *mCherry* transcription, after which cells were analysed by flow cytometry. (D) Percentage of mCherry-HI STC receiver cells and (E) median mCherry expression in STC receiver cells following 24 hours co-culture with CmGP1 sender cells (9:1 sender:receiver cell ratio), and doxycycline treatment for the indicated amount of time. Data presented as mean ± standard deviation of three independent experiments. A minimum of 7500 cells were analysed for each sample. (F) Distribution of mCherry fluorescence in STC receiver cells following co-culture with CmGP1 sender cells for 24 hours (9:1 sender:receiver cell ratio) and doxycycline treatment for the indicated amount of time. Data from a single experiment, representative of three biological replicates. STC clone A1, B1 and B2 cells cultured alone ("No senders") are displayed as shaded black histograms in all panels. 9000 cells were analysed for each sample. (G) Quantification of live imaging: median mCherry fluorescence intensity in STC clone A1, B1, B2 receiver cells and PSNBB-E control cells following 24 hours of co-culture with CmGP1GH1 (9:1 sender:receiver ratio) prior to treatment with 1μg/ml doxycycline. mCherry intensity was calculated at hourly timepoints. Movies for 0-24h and 24-48h periods were acquired separately; in order to display data on the same graph, the 24-48h data were scaled linearly, so that the 24 hour timepoint would match the 24 hour timepoint in the 0-24h movie. Scaling was performed independently for each of the four cell lines presented. Average of 3 biological replicates, 10 fields of view/replicate, minimum of 173 cells/replicate/timepoint. Error bars: standard deviation. A.F.U.: arbitrary fluorescence units.

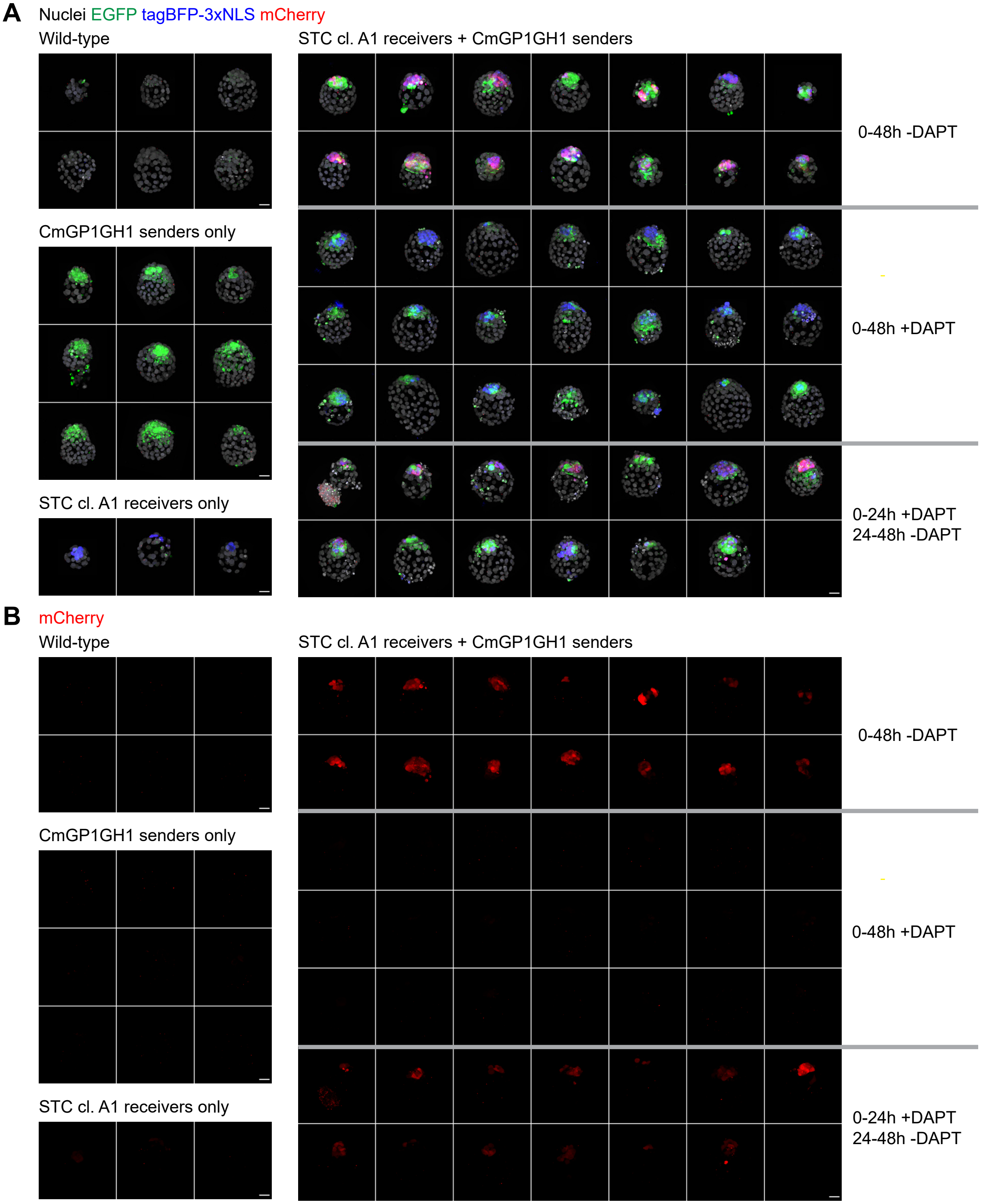


**Figure S17. DAPT treatment of chimaeric blastocysts.**
(A) Comparison of expression levels of EGFP, tagBFP-3xNLS and mCherry in wild-type and chimaeric blastocysts containing STC clone A1 and/or CmGP1GH1 sender cells. Following morula aggregation, embryos were either left untreated for 48 hours, treated with 100μM DAPT for 48 hours, or treated with 100μM DAPT for 24 hours, then washed and transferred to DAPT-free culture medium. Nuclei were counterstained with DRAQ7. Scale bar: 30μm. (B) Same embryos as shown in panel (A), displaying the mCherry channel only. Scale bar: 30μm. All images were acquired on a PerkinElmer Opera Phenix Plus.

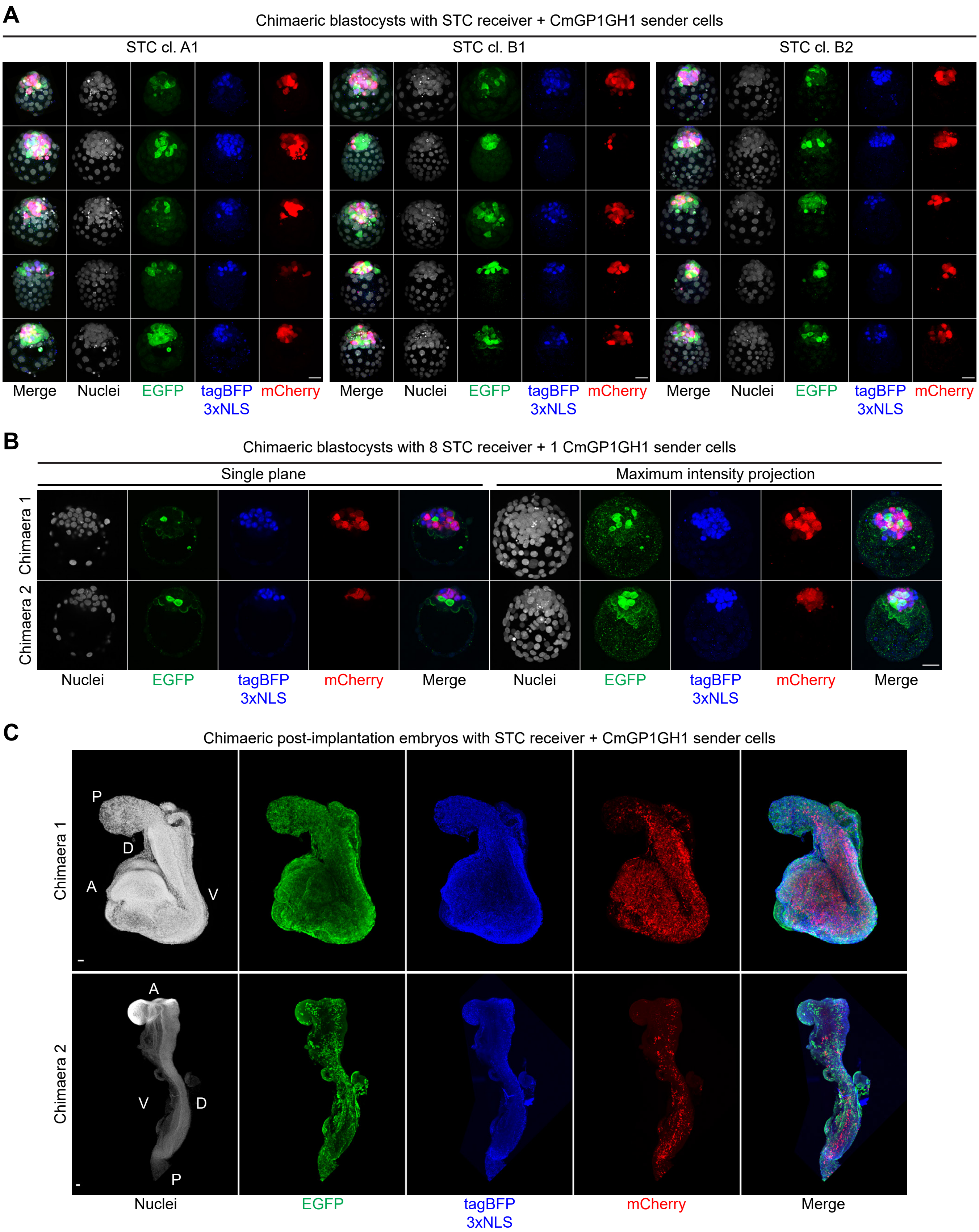


**Figure S18. SyNPL cells contribute to pre-implantation and post-implantation chimaeras.**
(A) Maximum intensity projections of chimaeric blastocysts containing STC clone A1, B1 or B2 receiver cells and CmGP1GH1 sender cells. Images were taken separately, and fluorescence intensities are not directly comparable. Nuclei were counterstained with DRAQ7. Scale bars: 30µm. (B) Chimaeric blastocysts generated by morula aggregation of 1 CmGP1GH1 and 8 STC clone A1 receiver cells. Nuclei were counter-stained with DRAQ7. Scale bars: 30µm. (C) Maximum intensity projections of post-implantation chimaeric embryos containing STC clone A1 and CmGP1GH1 sender cells. Images were taken separately, and fluorescence intensities are not directly comparable. Nuclei were counter-stained with DAPI (top) or DRAQ7 (bottom). Scale bars: 30µm. A: Anterior, P: Posterior, D: Dorsal; V: Ventral. Images in (A) and (C) were acquired on a Leica SP8, images in (B) were acquired on a PerkinElmer Opera Phenix Plus.

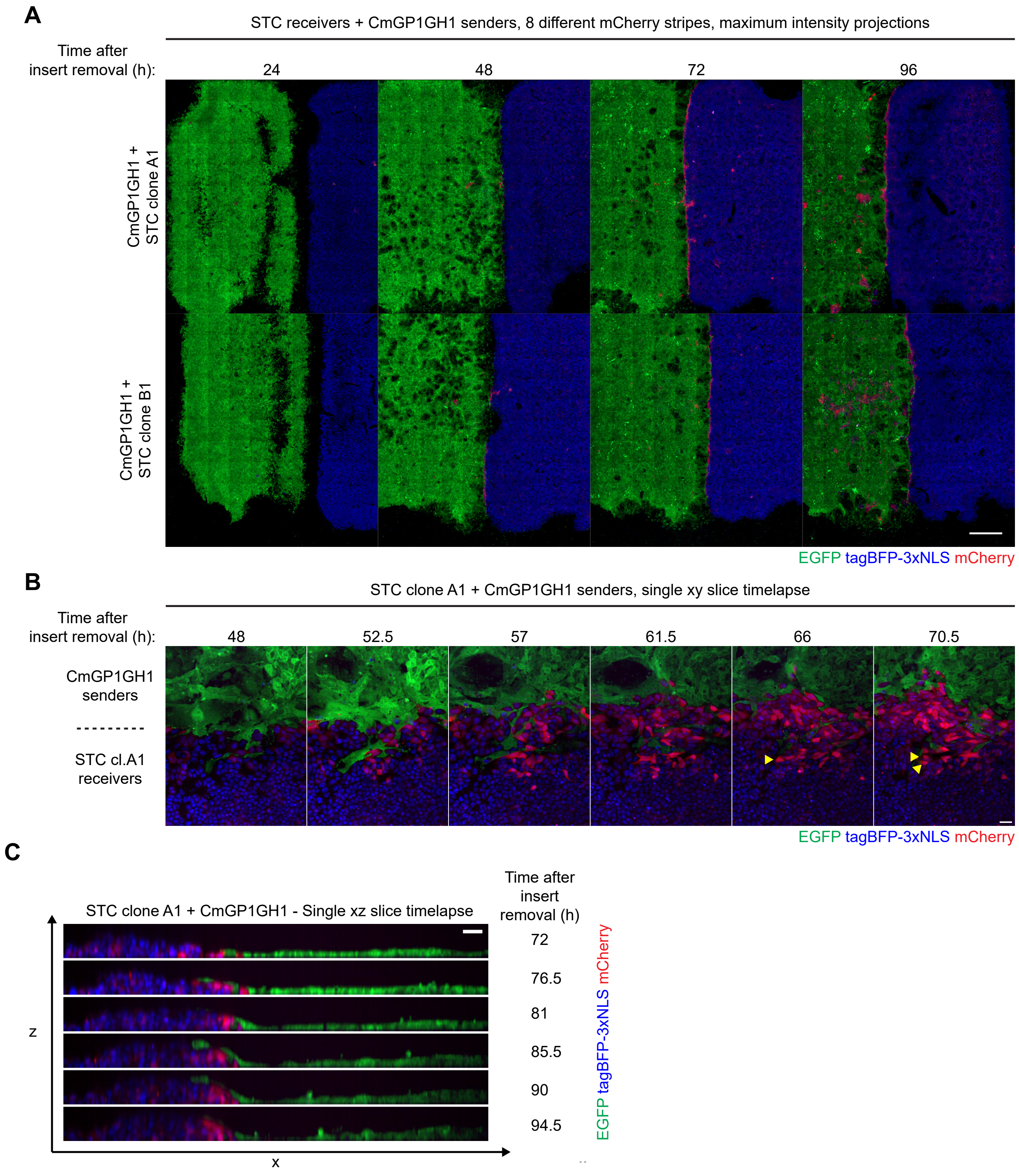


**Figure S19. Characterisation of synthetic stripe pattern of mCherry expression.**
(A) Low magnification view of synthetic stripe patterns of mCherry expression, generated by co-culture of STC clones A1 or B1 receiver cells with CmGP1GH1 sender cells. Each image represents a separate stripe. Scale bar: 1mm. (B) Timelapse imaging of border between STC clone A1 receiver and CmGP1GH1 sender cells. Photobleaching led to reduction in signal intensity over time. Single confocal slice at the bottom of the dish. Yellow arrowheads indicate activated receiver cell division in plane perpendicular to sender:receiver border, contributing to stripe diffusion.(C) Timelapse imaging of border between STC clone A1 receiver and CmGFP1GH1 sender cells. Photobleaching led to reduction in signal intensity over time. Single xz slice, showing thickness of border region. Sender cells can be seen migrating on top of receiver cells. Scale bar: 30µm. All images were acquired on a PerkinElmer Opera Phenix Plus microscope.

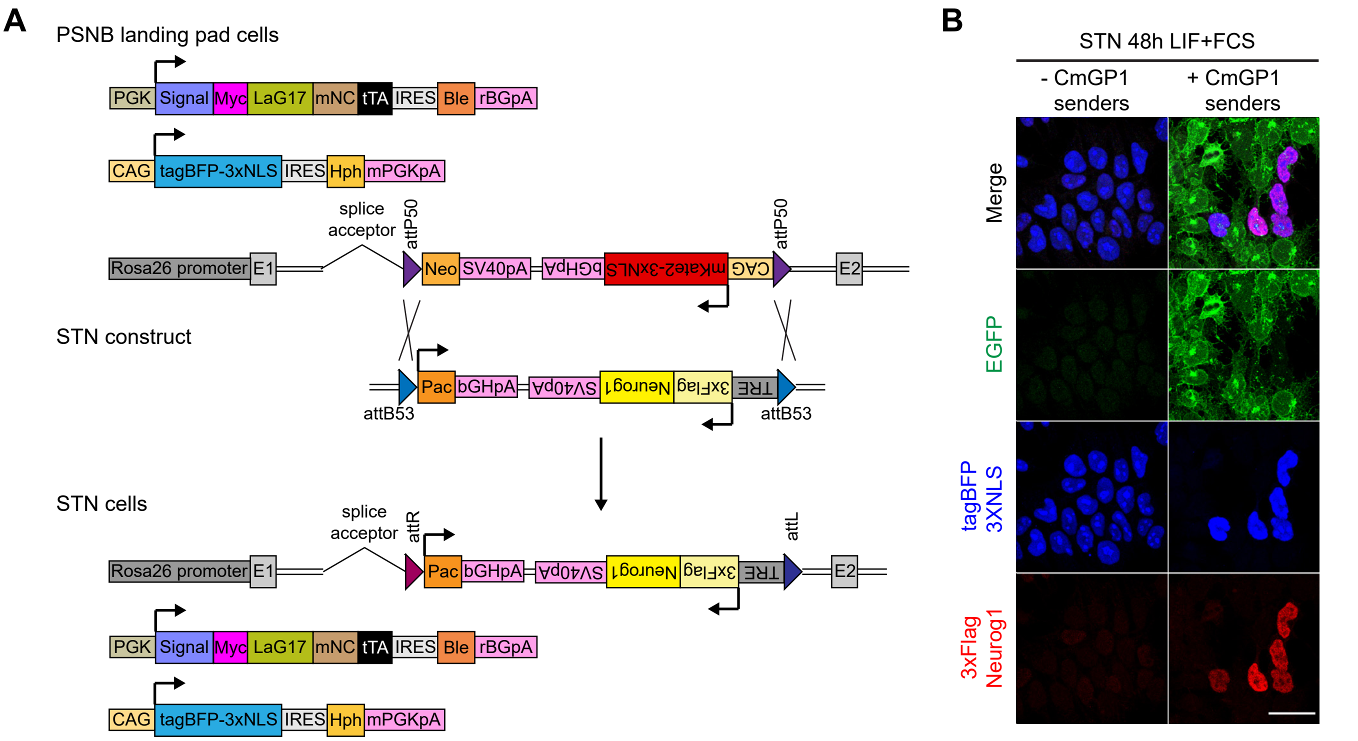


**Figure S20. Generation of Neurogenin1-inducible STN receiver cells.**
(A) Strategy to replace Neo-mKate2 cassette with STN cassette in PSNB landing pad ESCs through φC31 integrase-mediated RMCE. (B) EGFP, tagBFP and Flag immunofluorescence of STN receiver ESCs co-cultured for 48 hours with CmGP1 sender ESCs (9:1 sender:receiver cell ratio) in ESC culture medium ("LIF+FCS"). Scale bar: 30μm.

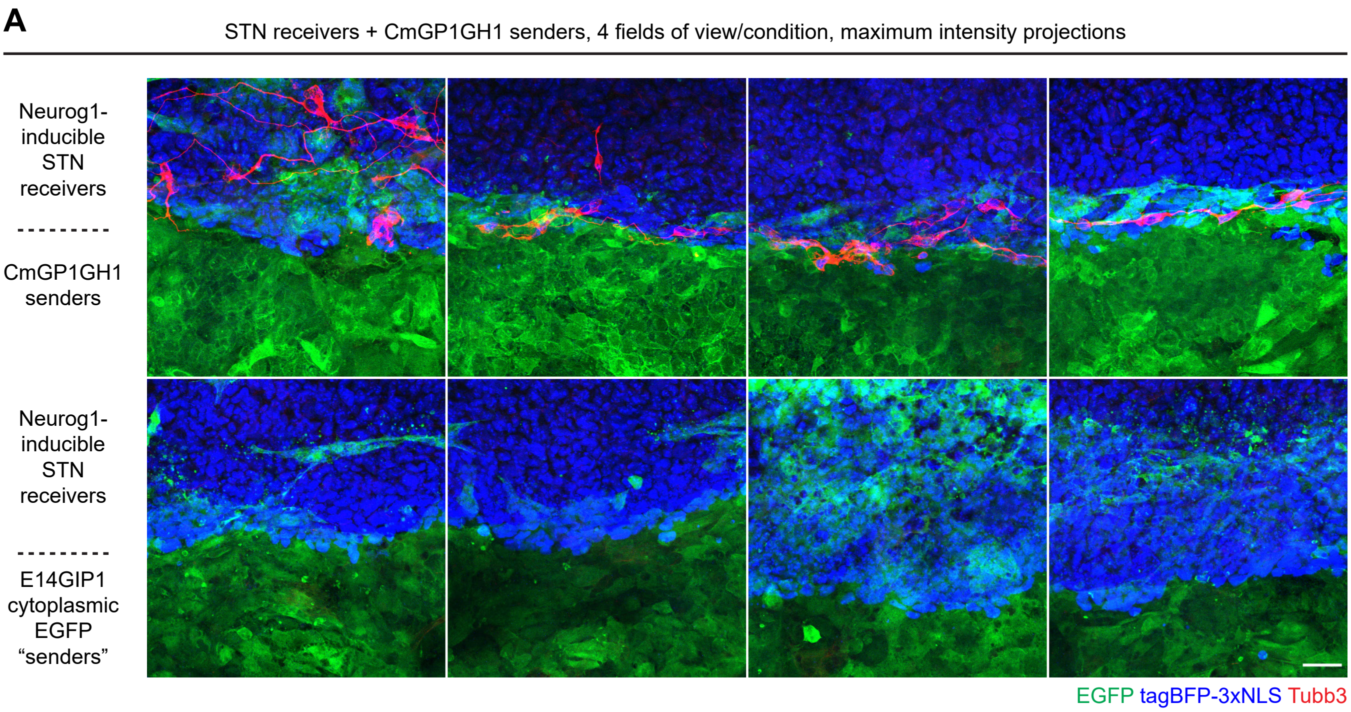


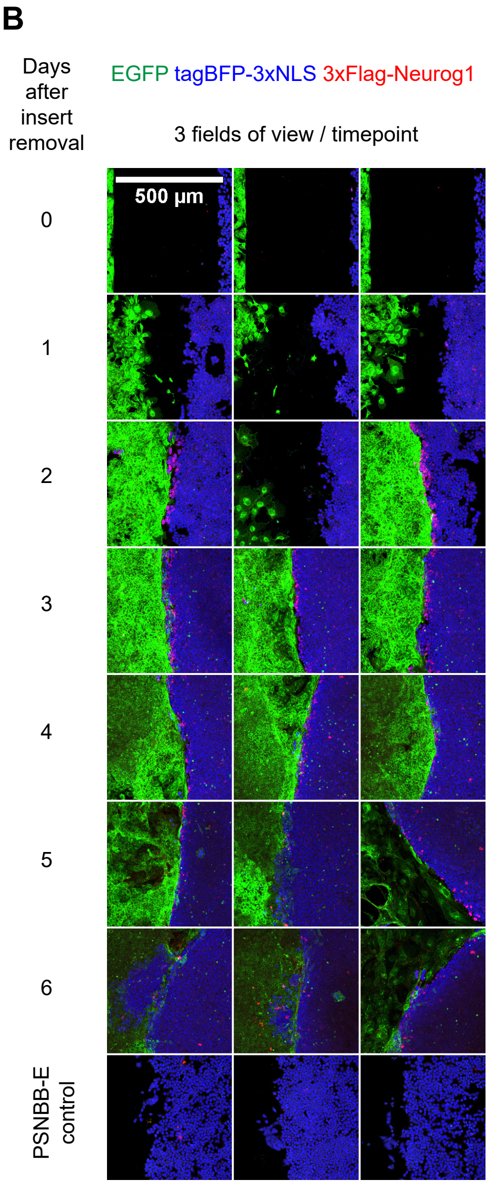


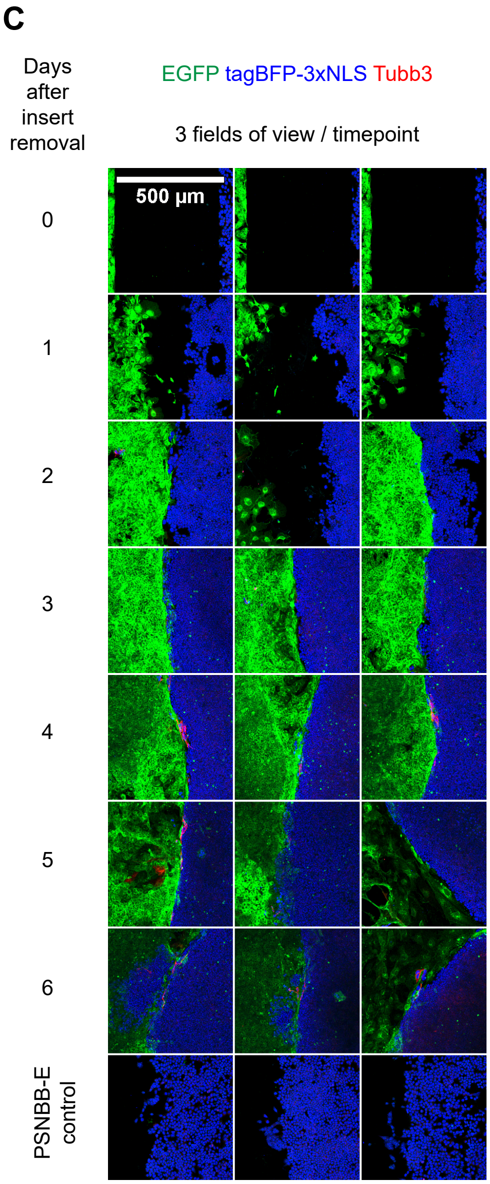


**Figure S21. Characterisation of neuronal differentiation stripe.**
(A) Immunofluorescence analysis of STN receiver cells co-cultured with either CmGP1GH1 sender cells or with E14GIP1 cytoplasmic EGFP-expressing control cells, illustrating that interaction with extracellular EGFP is required to drive neuronal differentiation in STN receiver cells. 4 fields of view are displayed per condition. Scale bar: 30µm.
(B-C) Immunofluorescence analysis of kinetics of (B) 3xFlag-Neurog1 or (C) Tubb3 induction. Maximum intensity projections of 3 fields of view at low magnification are displayed to capture variability. Scale bar: 500µm.

### Movie legends



**Movie 1. Time-lapse imaging of interactions between STC receiver and CmGP1GH1 sender cells.**
Time-lapse imaging of STC clone A1 receiver cells cultured with CmGP1GH1 sender cells at a 1:1 ratio for 24 hours. Brightfield (grey), EGFP (green) and mCherry (red) images taken every 10 minutes over a 24 hour period. Immunofluorescence analysis of tagBFP (blue), EGFP (green) and mCherry (red) expression at the 24 hour timepoint is shown at the end of the movie. All images were acquired using a Nikon Ti-E microscope.

**Movie 2. Kinetics of mCherry induction in STC receiver cells.**
Time-lapse imaging of STC clone A1 receiver cells cultured with CmGP1GH1 sender cells at a 1:9 ratio at high density for 24 hours. tagBFP-3xNLS (blue), GFP (green) and mCherry (red) images taken every hour over a 24 hour period. Imaging was performed on a PerkinElmer Opera Phenix Plus microscope.

**Movie 3. Time-lapse imaging of transient interactions between STC receiver and CmGP1GH1 sender cells.**
Time-lapse imaging of STC clone A1 receiver cells cultured with CmGP1GH1 sender cells at a 1:1 ratio for 24 hours. Brightfield (grey), EGFP (green) and mCherry (red) images taken every 10 minutes over a 24 hour period. Immunofluorescence analysis of tagBFP (blue), EGFP (green) and mCherry (red) expression at the 24 hour timepoint is shown at the end of the movie. All images were acquired using a Nikon Ti-E microscope. Movies 1 and 3 display different fields of view from the same experiment.

**Movie 4. Kinetics of mCherry downregulation in STC receiver cells (0-24h).**
Timelapse imaging of STC clone A1 receiver cells cultured with CmGP1GH1 sender cells at a 1:9 ratio for 24 hours, then treated with 1µg/ml doxycycline (to inhibit mCherry transcription) and imaged for 24 hours. Brightfield (grey), EGFP (green) and mCherry (red) images taken every 10 minutes over a 24 hour period, using a Nikon Ti-E microscope.

**Movie 5. Kinetics of mCherry downregulation in STC receiver cells (24-48h).**
Timelapse imaging of STC clone A1 receiver cells cultured with CmGP1GH1 sender cells at a 1:9 ratio for 24 hours, then treated with 1µg/ml doxycycline (to inhibit mCherry transcription) and imaged between 24 and 48 hours following addition of doxycyline. Brightfield (grey), EGFP (green) and mCherry (red) images taken every 10 minutes over a 24 hour period, using a Nikon Ti-E microscope.

**Movie 6. Time-lapse imaging of morula to blastocyst transition following morula aggregation of STC receiver and CmGP1GH1 sender cells.**
Time-lapse imaging of two wild-type morulae aggregated with STC clone A1 receiver cells and CmGP1GH1 sender cells. Brightfield (grey), EGFP (green) and mCherry (red) images taken every 30 minutes over a 27 hour 15 minute period. tagBFP-3xNLS (blue) was acquired for the first timepoint only, as sustained exposure to ultraviolet light results in death of chimaeric morulae. Filming started approximately 4 hours after morula aggregation.

As Movies 1, 3, 4 and 5 were acquired simultaneously and batch-processed, fluorescence intensities are directly comparable across these movies.
